# A comprehensive evaluation of taxonomic classifiers in marine vertebrate eDNA studies

**DOI:** 10.1101/2024.02.15.580601

**Authors:** Philipp E. Bayer, Adam Bennett, Georgia Nester, Shannon Corrigan, Eric J. Raes, Allison S. McInnes, Madalyn Cooper, Marcelle E. Ayad, Philip McVey, Anya Kardailsky, Jessica Pearce, Matthew W. Fraser, Priscila Goncalves, Stephen Burnell, Sebastian Rauschert

**Affiliations:** Minderoo Foundation, Perth 6000, WA; The UWA Oceans Institute, The University of Western Australia, Crawley 6009, WA; School of Biological Sciences, The University of Western Australia, Crawley 6009, WA

**Keywords:** Environmental DNA, Taxonomic classifiers, Machine Learning, Sequence alignment, Marine Biodiversity

## Abstract

Environmental DNA (eDNA) metabarcoding is a widely used tool for surveying marine vertebrate biodiversity. To this end, many computational tools have been released and a plethora of bioinformatic approaches are used for eDNA-based community composition analysis. Simulation studies and careful evaluation of taxonomic classifiers are essential to establish reliable benchmarks to improve accuracy and reproducibility of eDNA-based findings.

Here we present a comprehensive evaluation of nine taxonomic classifiers exploring three widely used mitochondrial markers (12S rDNA, 16S rDNA, and COI) in Australian marine vertebrates. Curated reference databases and exclusion database tests were used to simulate diverse species compositions, including three positive control and two negative control datasets. Using these simulated datasets, we were able to identify between 19% to 85% of marine vertebrate species using mitochondrial markers. We show that MMSeqs2 and Metabuli generally outperform BLAST with 10% and 11% higher F1 scores for 12S and 16S rDNA markers, respectively, and that Naive Bayes Classifiers such as Mothur outperform sequence-based classifiers except MMSeqs2 for COI markers by 11%. Database exclusion tests reveal that MMSeqs2 and BLAST are less susceptible to false positives compared to Kraken2 with default parameters. Based on these findings, we recommend that MMSeqs2 is used for taxonomic classification of marine vertebrates given its ability to improve species-level assignments while reducing the number of false positives. Our work contributes to the establishment of best practices in eDNA-based biodiversity analysis to ultimately increase the reliability of this monitoring tool in the context of marine vertebrate conservation.

## Introduction

Environmental DNA (eDNA) monitoring is one of the fastest growing biomonitoring tools, promising to revolutionise biodiversity measurements (Deiner et al., 2017, Cristescu and Hebert, 2018, Takahashi et al., 2023). eDNA-based studies tend to detect more species than conventional surveys (e.g., capture, visual or acoustic census) (Fediajevaite et al., 2021, Alexander et al., 2022, Nester et al., 2023), while diversity estimates and species inventories obtained by eDNA methods are generally congruent with conventional methods (Keck et al., 2022). For monitoring of marine vertebrate species, eDNA is particularly promising as conventional survey methods are often more resource- and time-intensive and lack scalability (Valentini et al., 2016, Beng and Corlett, 2020, Bessey et al., 2021).

Over the past decade, there has been a remarkable surge in the use of eDNA in biodiversity studies (Takahashi et al., 2023), positioning its application as a biomonitoring tool as a developing field in a ‘transitional phase’ (Schenekar, 2023). Consequently, there is a pressing need to establish benchmarks for generating accurate and reliable data and to identify the limitations of eDNA-based biodiversity survey approaches. Standards and guidelines are currently in development across the eDNA metabarcoding workflow for field and laboratory work (Pawlowski et al., 2020, Minamoto et al., 2021, De Brauwer et al., 2023, Samuel et al., 2021). However, to the best of our knowledge, no such effort has been undertaken for eDNA bioinformatic analyses.

The lack of bioinformatics standards may stem from a lack of consensus around computational methods and parameters, with most laboratories relying on bespoke analytical pipelines and/or customised databases unavailable to the public (Mousavi-Derazmahalleh et al., 2021, Takahashi et al., 2023). Bioinformatic parameters and thresholds which have been demonstrated to influence eDNA data (Alberdi et al., 2018, Mathon et al., 2021, Pearce et al., 2023) are often not reported, posing challenges to accurate taxonomic and ecological interpretations of the sampling data. A comprehensive evaluation of eDNA bioinformatics pipelines and tasks is crucial to promote consensus and reproducibility among researchers and increase confidence of management and government bodies in using eDNA data for decision making. As a step toward this, we introduced a standardised bioinformatics pipeline for marine vertebrate eDNA studies, assessing the impact of sequence read denoising settings on biological outcomes (Pearce et al., 2023). Additionally, we have begun submitting additional features such as VSEARCH post-clustering and phyloseq object creation to the *nf-core* pipeline *ampliseq* (Straub et al., 2020) and implemented the eDNA simulation steps presented in the current study in a *nf-core* pipeline *readsimulator* (https://github.com/nf-core/readsimulator).

A notable scarcity of eDNA-based benchmarks for marine vertebrates exists (Mathon et al., 2021), with few studies comparing taxonomic classifiers against known data. Bourret et al. (2023) tested different BLAST parameters using a regional curated subset of Metazoans of the BOLD database, revealing that database curation increased the number of species detected. To our knowledge, Mathon et al. (2021) conducted the sole marine vertebrate study available to date using simulated communities where they compared 13 bioinformatic programs and pipelines and recommended VSEARCH for taxonomic assignments. Only one study has carried out a meta-analysis of taxonomic classifiers to date, but all studies included focused on bacteria (Gardner et al., 2019). In that meta-analysis, k-mer-based tools such as Kraken2 and an implementation of the probabilistic classifier Naive Bayes Classifiers (NBC) showed an improved performance (Wang et al., 2007).

However, very few studies have utilised simulated data as recommended by Gardner et al. (2019), despite the widespread practice of conducting benchmark studies in other fields such as machine learning (Thiyagalingam et al., 2022). Benchmarks are crucial for accurately assessing the strengths and limitations of eDNA studies, as simulated data has the potential to address questions that real data may be unable to answer (O’Rourke et al., 2020, Lotterhos et al., 2022). In real data, where real species composition is unknown, it is not possible to assess classifier performance. Only simulated data, where species composition is known, can answer questions on accuracy of taxonomic classification.

For eDNA tools to be widely accepted, trusted, and routinely employed by management and government stakeholders, eDNA-based taxonomic profiles need to be reliable. To address this need, we present a comprehensive simulation-based study that evaluates the impact of taxonomic classifiers and underlying databases in marine vertebrate eDNA studies. We employed curated 12S rDNA, 16S rDNA, and COI reference databases including Australian species of marine mammals, reptiles, fish, and birds to simulate three highly diverse communities which were used to evaluate the performance of nine taxonomic classifiers. Furthermore, we designed an exclusion database to measure false positive rates in classifiers, integrated newly released classification software that was not available in previous comparative studies, and introduced customised machine learning-based approaches into our analysis. Our findings have the potential to contribute significantly towards the standardised practices and improved accuracy and confidence in eDNA-based biodiversity surveys. These advancements are pivotal in unlocking the full potential of eDNA, allowing it to progress from a ‘transitional’ (Schenekar, 2023) to a widely-adopted and trusted technology.

## Materials and Methods

### Custom databases of mitochondrial markers for Australian marine vertebrates

Custom curated reference databases have been shown to increase reliability and species-level detections in eDNA studies (Collins et al., 2021, Bourret et al., 2023, Jeunen et al., 2023). We built gene-specific reference databases for 12S rDNA, 16S rDNA, and COI including marine vertebrates that inhabit the Australian Exclusive Economic Zone (EEZ). Due to the lack of mitochondrial gene sequences available in public databases for species occurring within the Australian EEZ, reference databases were created at the family level to maximise the genetic diversity of included species. (Ahyong, 2023) (Table S1). For each family of marine vertebrates present in Australian waters (list sourced from FishBase and Australian Faunal Directory), all full and partial gene sequences for 12S, 16S, and COI as well as full mitochondrial genomes were downloaded from NCBI for all species within that family (including those that are not native to Australian waters).

For each species, family-level NCBI taxonomy IDs were extracted using taxonkit v0.12 *lineage* and *name2taxid* (Shen and Ren, 2021). All three mitochondrial genes and full mitochondrial genomes were downloaded for these taxonomy IDs using Entrez direct e-utilities v16.2 (full search terms in Supplementary Note 1), removing species with unspecific species-level labels (*cf., sp.*). For each of the marker genes, we searched for mislabelled species by self-blasting the downloaded sequences (e-value 1e-10) and calculating the LCA for each sequence using taxonkit *lca*, only including BLAST hits with a sequence identity above 97% and a query coverage of 100%. Gene queries with a high taxonomic level assigned (i.e., above the family level) indicates that at least one of the subject genes in the database is mislabelled. For each of these per-LCA sets of subjects, we counted the number of families using taxonkit *lineage* and labelled the family with fewer hits as potentially mislabelled and removed the corresponding subject-genes from the database.

### Simulating amplicon sequencing libraries

We opted to evaluate classifiers using synthetic datasets as it provides a known composition of species, referred to here as **positive control** datasets as outlined in Gardner et al. (2019). The synthetic datasets were generated by simulating the PCR amplification process using CRABS v0.1.1 (Jeunen et al., 2023) *insilico_pcr* allowing an error rate of 4.5 (--error 4.5) for the primers 12S_Miya and 16S_Berry, and 10 (--error 10) for COI_Leray. Amplicon-sequencing products were simulated by extracting the mitogenome regions for 12S_Miya (Miya et al., 2015), 16S_Berry (Berry et al., 2017), and COI_Leray (Leray et al., 2013) (Table S2). We chose the Miya, Berry, and Leray primers as representative primers for these three gene regions as they are most commonly used in marine eDNA studies (Collins et al., 2019, Miya, 2022).

Using these primer sets, we generated three simulated query datasets: one **‘balanced’** dataset containing highly diverged marine vertebrates based on families present in the Australian EEZ, one **‘similar’** dataset containing highly similar Actinopterygii (family Lutjanidae), and one **‘realistic’** dataset based on GBIF.org sightings data around Wadjemup (Rottnest Island, Western Australia). For the **‘balanced’** dataset, for 12S, 16S, and COI respectively we selected the most distinct and unique sequences by aligning the simulated PCR-products using MUSCLE v5.1.linux64 (Edgar, 2004) and inferring phylogenies using modeltest-ng v0.1.7 and raxml-ng v1.2.0 (Kozlov et al., 2019). From each phylogeny, we chose the 100 most-representative species that cover most of the diversity in the phylogeny using PARNAS v0.1.3 (option -n 100). This resulted in three distinct sets of chosen genes for 12S, 16S, and COI due to species-specific genetic diversity within these three marker genes. The chosen species are not necessarily Australian species, as we downloaded all sequences for taxonomic families present in the EEZ. The **‘similar’** dataset consists of 12S, 16S and COI genes of 36 publicly available species mitogenomes for fish within the family Lutjanidae (Table S3). The **‘realistic’** dataset was generated by downloading all Elasmobranchii and Actinopterygii species sighted around Wadjemup (Rottnest Island) from GBIF.org (accessed 01 August 2023, GBIF occurrence download https://doi.org/10.15468/dl.ynd3x7), and extracting all simulated 12S, 16S, and COI PCR products for the available mitogenomes. The resulting dataset contained mitogenomes for 302 out of the 387 sighted species (Table S4).

For each primer and for each of the ‘balanced’, ‘similar’, and ‘realistic’ datasets, five amplicon sequencing libraries were simulated using ART v2.5.8 *art_illumina*.(Huang et al., 2012). ART was chosen as, to our knowledge, it is the only sequencing simulator capable of simulating amplicon sequencing using realistic Illumina-sequencing-based error profiles. The chosen parameters were: -ss HS25 (simulate Illumina HiSeq 2500 error profile), -amp (amplicon sequencing), -f 20 (fold of 20 per species), and -l 130 (simulate 130 bp long reads, accounting for the usual 20 bp due to index and primer sequences in Illumina sequencing) and five different seeds (42 to 46).

Different read depths have an impact on biologically relevant measurements such as alpha- and beta-diversity (Shirazi et al., 2021). In fish eDNA studies, a read depth of 50,000 read pairs has been recommended per sample to adequately capture diversity measures (García-Machado et al., 2023). We therefore simulated 500 read pairs per gene-copy and species to ensure sufficient read-based representation for each gene-copy. We chose an equal number of reads per gene-copy and species as we are assessing taxonomic classifiers, not eDNA analysis pipelines, and read depth variation may confound the impact of classifiers with the impact of ASV analysis pipeline. The entire pipeline simulating amplicon sequencing reads at different depths per primer set was codified in a *nf-core* Nextflow pipeline and is available for future eDNA simulation studies with different primers, sequencing technologies, or species and datasets of interest (https://github.com/nf-core/readsimulator).

In taxonomic classifiers, employing negative controls (random sequences or sequences of distant relatives outside the study scope) is important to measure false positives as no negative control should have assigned taxonomic classification (Gardner et al., 2019). We generated two negative controls: One based on random sequences, and one based on bacterial sequences.

We generated 100 randomised gene sequences using a custom Python-script selecting 170 bp of A, T, G, and C to result in genes of a similar length to the 16S PCR product (*gene_faker.py*). This was done for each gene region (12S, 16S and COI) to simulate three negative control libraries with 500 reads per gene (the **random negative control**).

As random DNA sequences might not be recognisable by marker gene-specific classifiers, we also used bacterial and archaeal genes from Greengenes v13_5 (Table S5). For each of the 100 most common species in Greengenes, one gene was extracted at random for each gene region of interest (16S, 12S, COI). To generate products of similar length to the PCR products of the primers (Berry_16S, 12S_Miya, and COI_Leray), *in silico* PCR products were extracted from 164 to 191 bp (12S_Miya), 149bp to 240 bp (16S_Berry), and 310bp to 313 bp (COI_Leray). This resulted in 500 simulated amplicon reads per gene, using the OceanOmics-amplicon-nf pipeline to merge simulated reads into ASVs (the **microbial negative control**).

### Comparing different taxonomic classifiers

The negative and positive control eDNA libraries were merged into amplicon sequence variants (ASVs) using the Oceanomics-amplicon-nf pipeline release Thalassa (Pearce et al., 2023) https://github.com/MinderooFoundation/OceanOmics-amplicon-nf). We chose default settings in the pipeline except for COI, where we set the minimal overlap between R1 and R2 to 0 as the primer product COI_Leray is larger than the total length of R1 and R2. We have chosen nine taxonomic classifiers across different settings, ranging from standard BLAST+ blastn to modern k-mer-based approaches (Table 1). We used identical settings where possible: e-value cut-off of 1e-5, percentage identity cut-off of 97%, and a query coverage of 100%. Both the full and all exclusion databases were used as reference databases. We evaluated several parameters for some classifiers such as BLAST, MMSeqs2, and Kraken2. Due to a gap in the middle of the COI primer-product due to the simulated PCR product being longer than the total length of the forward and reverse read, alignments reported by both MMSeqs2 and VSEARCH were shorter than roughly 50%. Consequently, we lowered the query-percentage cutoff to 0.4 in both programs.

**Table 1:** Overview of the chosen taxonomic classifiers, their corresponding citation, chosen settings, and the version numbers.

| Tool name (shorthand) | Reasoning for inclusion | Settings | Version number | Citation |
| --- | --- | --- | --- | --- |
| **K-mer or alignment-based tools** |  |  |  |  |
| BLAST+ blastn with LCA (BLAST97) | Commonly used in eDNA studies, i.e. (Mousavi-Derazmahalleh et al., 2021) | -e 1e-5 (blastn), 97% identity, 100% query coverage (LCA script) | 2.13.0+ | (Camacho et al., 2009, Mousavi-Derazmahalleh et al., 2021) |
| (BLAST100) | More stringent identity cutoffs as argued for in bacteria (Edgar, 2018) | 100% identity, 100% query coverage |  |  |
| DADA2 assignSpecies (DADA2Spec) | Standard tool | tryRC = TRUE, allowMultiple = TRUE | 1.26.0 | (Callahan et al., 2016) |
| MMSeqs2 easy-taxonomy (MMSeqs2_97) | has been shown to be more sensitive than BLAST in some cases (Steinegger and Söding, 2017) | easy-taxonomy --cov-mode 2 -c 1 --min-seq-id 0.97 -e 1e-5 --search-type 3 --orf-filter 0 (for COI: -c 0.4) | 14.7e284 | (Steinegger and Söding, 2017) |
| (MMSeqs2_100) |  | --min-seq-id 1<br>(for COI: -c 0.4) |  |  |
| Metabuli | Incorporates amino-acid level comparisons for more distant hits | --seq-mode 3 --<br>accession-level 1 --<br>min-cons-cnt-euk 4 -<br>-min-cons-cnt 4 | branch<br>newScore,<br>commit<br>ba7375e | (Kim and Steinegger, 2023) |
| QIIME2 classify-consensus-vsearch (VSEARCH) | A QIIME2 wrapper around VSEARCH, commonly used in eDNA studies (i.e., (García-Machado et al., 2023, He et al., 2023, Feng et al., 2023)) | --p-perc-identity 0.97 --p-query-cov 1<br>(for COI: --p-query-cov 0.4) | QIIME2 2020.8<br>VSEARCH v2.7.0 | (Rognes et al., 2016) |
| Kraken2 (Kraken2_0.0) | shown to outperform DADA2 and QIIME2 in bacterial 16S amplicon sequencing mock communities (Odom et al., 2023), but shown to report false positives with incomplete reference databases (Gihawi et al., 2023) | defaults | 2.1.3 | (Wood et al., 2019) |
| (Kraken2_0.05) | Higher confidence cutoffs to reduce false positives | --confidence 0.05 |  |  |
| (Kraken2_0.1) |  | --confidence 0.1 |  |  |
| Naive Bayes Classifier-based tools |  |  |  |  |
| DADA2 assignTaxonomy (DADA2Tax) | Naive Bayes Classifiers have been shown to classify near the performance limit for 16S microbial genes classification (Ziemski et al., 2021) and across metagenomics classifier tasks (Wang et al., 2007, Gardner et al., 2019, Straub et al., 2020). | minBoot = 95, tryRC = TRUE | 1.26.0 | (Callahan et al., 2016) |
| Mothur |  | classify.seq with cutoff=97 | 1.48.0 | (Schloss et al., 2009, Schloss, 2020) |
| QIIME2 Naive Bayes Classifier (QIIME2) |  | Score cutoff of 0.97 | 2020.8 | (Bokulich et al., 2018) |
| Hyperparameter-optimised scikit-learn-based Naive Bayes Classifier (CustomNBC) | Naive Bayes Classifier hyperoptimised using scikit-learn's GridSearchCV | Class probability cutoff 0.97, hyperparameter optimisation via GridSearchCV (HashingVectorizer: ngram-range 8 to 46, MultinomialNB: alpha 0.001 to 0.1, fit_prior True to False). | 1.2.1 | (Pedregosa et al., 2011) |

Classification tasks are always a trade-off between recall (high true positives) and specificity (high true negatives). In eDNA studies, the level of specificity needed, and the consequences of false positives and false negatives on the objective and result, can vary significantly. For example, researchers conducting biodiversity studies typically prioritise detecting as many species as possible, including rare species, aiming for increased sensitivity while minimising false negatives. Conversely, in the context of monitoring invasive species, ensuring that the invasive species is not detected in areas where it has not yet established is crucial, calling for increased specificity and minimal to no false positives. These different outcomes are tracked using different measurements. For each classifier we measured true positives (TP), false positives (FP), true negatives (TN), and false negatives (FN). These four statistics were used to calculate recall (also known as sensitivity) via TP / TP + FN, specificity (true negatives) via TN / TN + FP, precision (false positives) via TP / TP + FP, the F1-score ((2 * precision * recall) / (precision + recall), and the F0.5-score (((1 + 0.5^2) * precision * recall) / (0.5^2 * precision + recall), i.e., F-score with a higher importance on precision).

### Different levels of clade exclusions

Carrying out classifier comparisons with sequenced reads from completely known species does not accurately represent real eDNA sequencing, where many of the species present in the environmental samples have not had their genomes sequenced (summarised in De Jong). One potential solution to simulate more realistic datasets is ‘clade exclusion’, where species are present in the sequenced samples that are not present in the reference database, or have been removed from the reference database (Peabody et al., 2015). We therefore generated three subsets of the 12S, 16S, and COI reference databases by excluding families. For each reference database, we randomly excluded entire families by removing all sequences belonging to 30%, 50%, or 70% of families in the database. The random choice of families to remove was repeated ten times each for a total of 30 exclusion reference databases.

The analyses presented in this paper are fully reproducible via targets (Landau, 2021) and hosted at https://github.com/MinderooFoundation/OceanOmics-classifier-comparison/.

## Results

### Custom databases of mitochondrial markers for Australian marine vertebrates

All available 12S, 16S, and COI fragments for Australian marine vertebrates (mammals, reptiles, fishes, and birds) were downloaded from NCBI on the 27^th^ of June 2023 using a curated list of species and GBIF.org-based sightings around Australia. Potential mislabels were automatically removed as curated reference databases generate more accurate species assignments (Collins et al., 2021).

The resulting curated databases contained 35,312 12S fragments, 34,139 16S fragments, and 171,539 COI fragments across 350, 344, and 357 families respectively. Within the three databases, most sequences were from Actinopterygii (ray-finned fishes) (12S 77%:, 16S: 79%, COI: 82%), followed by Reptilia (birds, reptiles, turtles) (12S: 15%, 16S: 16%, COI: 7%), then Chondrichthyes (cartilaginous fishes) (12S: 5%, 16S: 5%, COI: 11%), and finally Mammalia (mammals) (12S: 3%, 16S: 1%, COI: 1%). Among the most common Actinopterygii families were Gobiidae (12S: 9%, 16S: 6%, COI: 8%) and Serranidae (12S: 6%,16S: 5%, COI: 4%), two of the largest and most species diverse families. The databases are available at https://github.com/MinderooFoundation/OceanOmics-AmpliconReference.

### Simulating amplicon sequencing libraries

We simulated 12S_Miya (Miya et al., 2015), 16S_Berry (Berry et al., 2017), and COI_Leray (Leray et al., 2013) PCR products using all genes available for the species chosen for each positive and negative control dataset. The ASV simulation pipeline denoised a similar amount of ASVs as there were species in the input dataset (Table 2). In the ‘realistic’ Wadjemup dataset, there were more ASVs denoised than actual species owing to the presence of a diverse set of 141 *Phyllopteryx taeniolatus* (Common Seadragon) 12S, 16S, and COI genes in the reference databases. The program we chose to denoise reads into ASVs, DADA2, is designed to identify fine-scale variations within species (Callahan et al., 2016), resulting in a slightly higher number of sequences representing distinct populations of *P. taeniolatus* being denoised into separate ASVs. We also denoised these sequences using VSEARCH which resulted in a usually identical number of zOTUs (one less zOTU in Wadjemup, one more zOTU in Lutjanidae, identical in others) and decided to proceed with DADA2 ASVs only (Table S6).

**Table 2:** Composition of positive and negative control datasets.

| Type | Dataset | Marker | Genes | Species | Simulated ASVs |
| --- | --- | --- | --- | --- | --- |
| Positive, similar | Lutjanidae | 12S_Miya | 36 | 26 | 24 |
| Positive, similar | Lutjanidae | 16S_Berry | 36 | 26 | 27 |
| Positive, similar | Lutjanidae | COI_Leray | 36 | 26 | 27 |
| Positive, realistic | Wadjemup | 12S_Miya | 299 | 94 | 102 |
| Positive, realistic | Wadjemup | 16S_Berry | 302 | 97 | 112 |
| Positive, realistic | Wadjemup | COI_Leray | 302 | 97 | 117 |
| Positive, balanced | 100 Australian family marine vertebrates | 12S_Miya | 100 | 100 | 99 |
| Positive, balanced | 100 Australian family marine vertebrates | 16S_Berry | 100 | 100 | 99 |
| Positive, balanced | 100 Australian family marine vertebrates | COI_Leray | 100 | 100 | 99 |
| Negative | Greengenes, bacteria | 16S_Berry | 100 | 100 | 99 |
| Negative | Random | N/A | 100 | 100 | 100 |

### Comparing different taxonomic classifiers

On average, classifiers correctly assigned species labels to 48% of ASVs. Classifier performance varied widely, with the lowest assignment of 1% (1 out of 99 ASVs) and 0% observed using VSEARCH and DADA2Spec on the COI_Leray ‘balanced’ dataset, and the highest at 85% (23 out of 27 ASVs) in the ‘similar’ COI_Leray dataset using CustomNBC, MMSeqs2_97, and Mothur (Figure 1). We observed strong differences in classification accuracy between the 12S_Miya, 16S_Berry, and COI_Leray markers within the ‘similar’ Lutjanidae dataset. In COI_Leray, classifiers demonstrated higher accuracy assigning up to 85% of species correctly (CustomNBC, MMSeqs2_97, Mothur, Qiime2), followed by the 12S_Miya-based classifiers (Metabuli) which assigned up to 71% of species correctly. In contrast, 16S_Berry-based classifiers exhibited lower accuracy assigning up to 41% of species correctly (Metabuli). This discrepancy in classifier performance and accuracy between the three genes can be explained by the differing variability of the gene marker, with the 16S gene being highly similar or identical across ‘similar’ Lutjanidae species.

**Figure 1:**
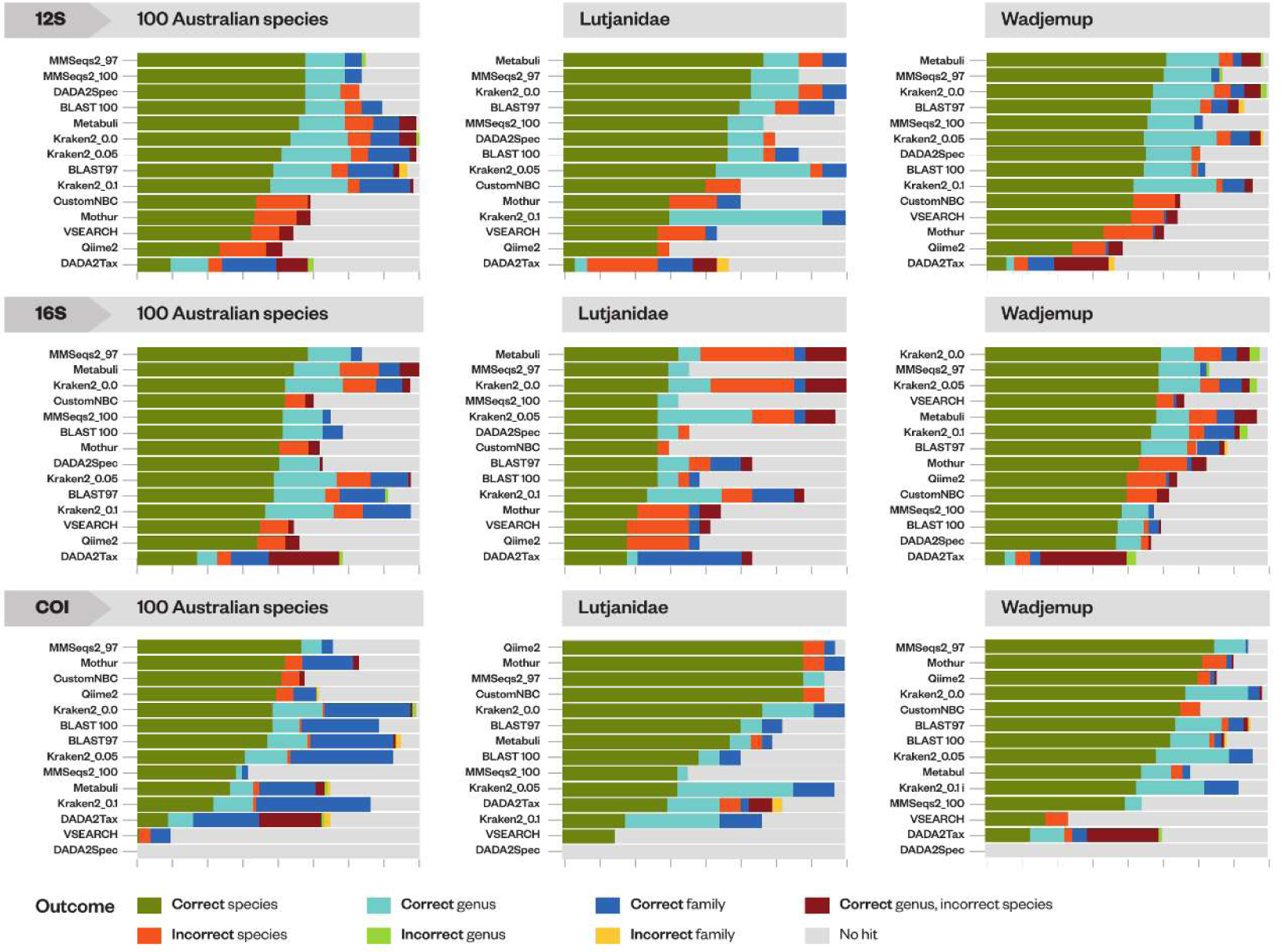
Percentages of correct and incorrect classifications at various levels of the biological hierarchy (species, genus, and family) for three simulated query datasets for 12S, 16S, and COI. For each classifier, species, genus, and family predictions were compared with the truth for simulated ASV sequences. Classifiers are sorted by the percentage of correct species.MMSeqs2_97 and MMSeqs2_100 denote percentage cut-offs, MMseqs2 with a 97% and 100% identity cutoff. BLAST97, BLAST100: BLAST with a 97% and 100% identity cutoff. Kraken2_0.0, Kraken2_0.05, Kraken2_0.1: Kraken2 with confidence cutoffs of 0 (default), 5%, and 10%.

Using the default values, Kraken2 often exhibits overconfidence in assigning species labels. For example, in the 16S_Berry ‘similar’ Lutjanidae dataset Kraken2 assigns species-level labels to 22 out of 27 ASVs, of which 12 ASVs (55%) were mislabelled. Adjusting Kraken2 confidence cutoffs to 0.05 in the ‘similar’ Lutjanidae dataset reduced the number of false positives, however this adjustment slightly lowered the number of correct species labels, with 7 out of 27 ASVs (26%) mislabelled.

When classifiers were wrong, they generally predicted the correct genus but the wrong species (Figure S1, Table S7). For example, using default cutoffs Kraken2 correctly identified the genus and species for 8 out of 22 ASVs (36%) for ‘similar’ Lutjanidae with 16S_Berry, and predicted the wrong genus and wrong species for just 4 out of 22 ASVs (18%).

We calculated accuracy, precision, recall, F1 and F-0.5-scores to evaluate true and false positives across all taxonomic classifiers (Figure 2, Table S8-S16). In 12S_Miya and 16S_Berry, Metabuli and MMseqs2 generally had the same highest accuracy scores of 0.67 and 0.66 for 12S and 0.56 for 16S_Berry compared with the median accuracy across all tools of 0.58 and 0.47 for 12S and 16S_Berry respectively.

**Figure 2:**
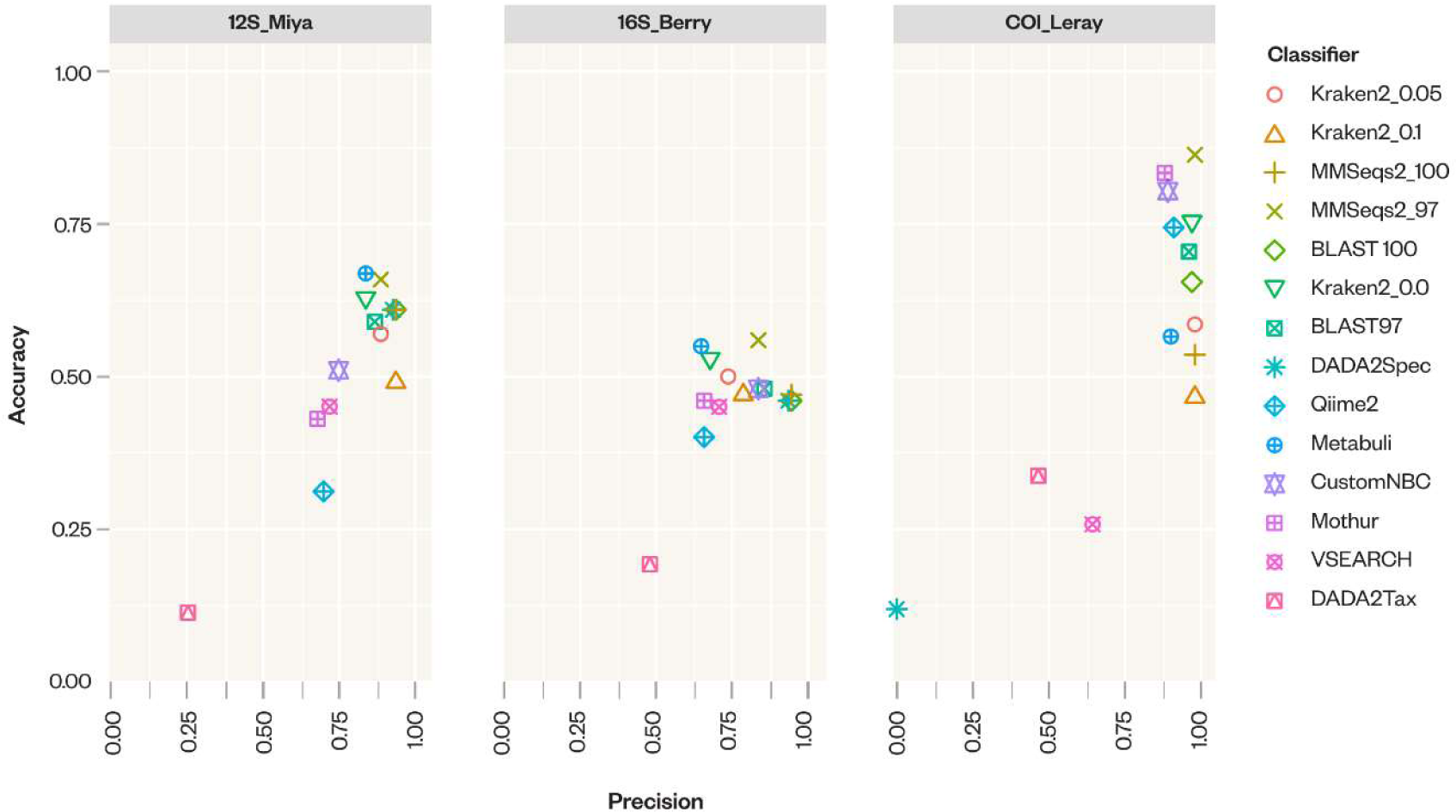
Precision compared with accuracy of taxonomic classifiers across 12S_Miya, 16S_Berry, COI_Leray based on medians across all positive control datasets.

In COI_Leray, Mothur had the highest accuracy with 0.84, 33% higher than the median accuracy of 0.63. In 12S_Miya, Kraken2_0.1 and MMseqs2_100 had the highest precision followed by DAD2Spec (0.94 and 0.93, all tools median: 0.86), while in 16S_Berry, MMSeqs2_100 and BLAST_100 had the highest precision (0.95 in both tools, all tools median: 0.77). In COI_Leray, on the other hand, Kraken2_0.05, Kraken2_0.1, MMseqs_97, and MMSeqs2_100 had the highest precision (0.99 in all cases) compared with the median precision of 0.95.

F1 and F0.5 scores display a similar trend, with Metabuli having the highest F1 scores in 12S_Miya (0.8, all tool median 0.72) and MMseqs2_97 having the highest F1 score in 16S_Berry (0.7, all tool median 0.62) and COI_Leray (0.92, all tool median 0.73). F0.5 scores are highest in 12S_Miya for BLAST100 and MMSeqs2_100 (0.85 for both, all tool median: 0.81) and in 16S_Berry, as well (0.78 for both tools, all tool median of 0.71). In COI_Leray, MMseqs2_97 had the highest F0.5 score as well (0.96, all tool median0.86). However, in COI_Leray, Naive Bayes-based tools Mothur, CustomNBC, and Qiime2 had the second to fourth highest F1-scores after MMSeqs2 (0.89, 0.87, and 0.84 respectively). DADA2Spec showed no hits in COI_Leray.

The assignment of incorrect or false positive species-labels can have negative repercussions for ecological assessments. For example, in the balanced query dataset (16S_Berry), ASV_53 is labelled as *Anguilla reinhardtii* (Australian Longfin Eel), a species listed as least concern (Pike et al., 2019b). However, VSEARCH mislabels this ASV as *A. australis* (Shortfin Eel) which is listed as near threatened (Pike et al., 2019a) (Table S17). Similarly, ASV_74 in the balanced dataset (12S_Miya) is *Lepidochelys kempii* (Kemp’s Ridley) a critically endangered species (Wibbels and Bevan, 2019). However, BLAST100 does not assign a species-level label to this ASV, while BLAST97 and Kraken2 assign the label *L. olivacea* (Olive Ridley Turtle) a species listed as vulnerable (Abreu-Grobois and Plotkin, 2008). In cases where classifiers falsely identify species of less concern than the endangered species, measures of protection based on eDNA assessment may fall short of what is required to establish effective species protection.

Conversely, classifiers falsely identifying species that are endangered or critically endangered instead of a species of lessor concern can have implications for the management of marine parks as protection status or resources may be incorrectly assigned or distributed. ASV_112 in the realistic dataset (16S_Berry) had no species-level label as there were too many carcharhinids with identical16S_Berry sequences. However, across all confidence levels Kraken2 and MMSeqs2 labelled this ASV as the endangered *Carcharhinus plumbeus* (Sandbar Shark) (Rigby et al., 2021).

Additionally, these errors can impact commercial evaluations. For example, ASV_67 in the 12S realistic Wadjemup dataset is *Scomber australasicus* (blue mackerel) and could not be labelled at the species level by BLAST, Kraken2, or MMSeqs2. While Mothur, CustomNBC, and Qiime2 labelled it as *S. colias* (Atlantic chub mackerel) and VSEARCH as the commercially important *S. japonicus* (Pacific chub mackerel; (Scoles et al., 1998). Such misidentifications can lead to incorrect decisions when establishing or assessing fish stocks of commercial value through use of eDNA-derived data. We next assessed the classifier performance with the negative control datasets with the expectation that no classifiers will find any marine vertebrates in the bacterial 16S genes (Greengenes) or the random DNA. As expected, no classifier assigned any species to the Greengenes or the random DNA dataset using the reference databases as these databases contain only marine vertebrates.

### Different levels of clade exclusions

In real or non-simulated scenarios, reference databases almost never include all reference sequences. To test the impact of incomplete reference databases on species assignments, all taxonomic classifiers were used to assign taxonomic labels to the five query datasets using three different levels of exclusion databases: 30% of families randomly removed, 50% of families randomly removed, and 70% of families randomly removed. Employing exclusion databases offers a more realistic way to assess the accuracy and performance of taxonomic classifiers in real-life settings. As exclusion databases have been made artificially incomplete, they offer a chance to measure over-confidence of classifiers: by removing all members of an ASV’s taxonomic family from the reference database we can measure whether a given classifier will mislabel this ASV or refuse to label it.

In the exclusion tests, different classifiers showed different levels of false positives. MMSeqs2_97 and BLAST100 (BLAST with a percentage identity cutoff of 100%) generally showed the highest F1 and F0.5 scores across all marker genes and percentages of randomly removed families in 12S_Miya, 16S_Berry, and COI_Leray simulated datasets (Table 3, Table S18).

**Table 3:** Averaged Accuracy, Precision, Recall, F1, and F0.5 scores for the highest F1-scoring classifier across 12S, 16S, COI, and the three types of exclusion database.

| Marker | Classifier with highest F1 score | Families removed | Accuracy | Precision | Recall | F1 | F0.5 |
| --- | --- | --- | --- | --- | --- | --- | --- |
| 12S_Miya | MMSeqs2_97 | 30% | 0.41 | 0.82 | 0.45 | 0.58 | 0.7 |
| 12S_Miya | BLAST100 | 50% | 0.3 | 0.9 | 0.31 | 0.47 | 0.65 |
| 12S_Miya | BLAST100 | 70% | 0.16 | 0.92 | 0.17 | 0.28 | 0.48 |
| 16S_Berry | MMSeqs2_97 | 30% | 0.44 | 0.93 | 0.46 | 0.61 | 0.77 |
| 16S_Berry | MMSeqs2_97 | 50% | 0.3 | 0.93 | 0.31 | 0.46 | 0.66 |
| 16S_Berry | MMSeqs2_97 | 70% | 0.17 | 0.89 | 0.18 | 0.3 | 0.49 |
| COI_Leray | MMSeqs2_97 | 30% | 0.43 | 0.87 | 0.46 | 0.6 | 0.74 |
| COI_Leray | MMSeqs2_97 | 50% | 0.31 | 0.85 | 0.33 | 0.48 | 0.65 |
| COI_Leray | MMSeqs2_97 | 70% | 0.16 | 0.87 | 0.16 | 0.28 | 0.47 |

Species diversity can be described by the number of detected species. We hypothesised that different classifiers would yield different species-level diversities especially across different levels of exclusion (Figure 3). We found strong discrepancies in false positives across classifiers when using exclusion databases in 12S_Miya, 16S_Berry, and COI_Leray. In general, Kraken2_0.0 resulted in significantly higher numbers of unique species counts (p < 0.05) than other classifiers due to many false positives. MMSeqs2 detected a slightly higher species level diversity than BLAST; however, this difference was not statistically significant.

**Figure 3:**
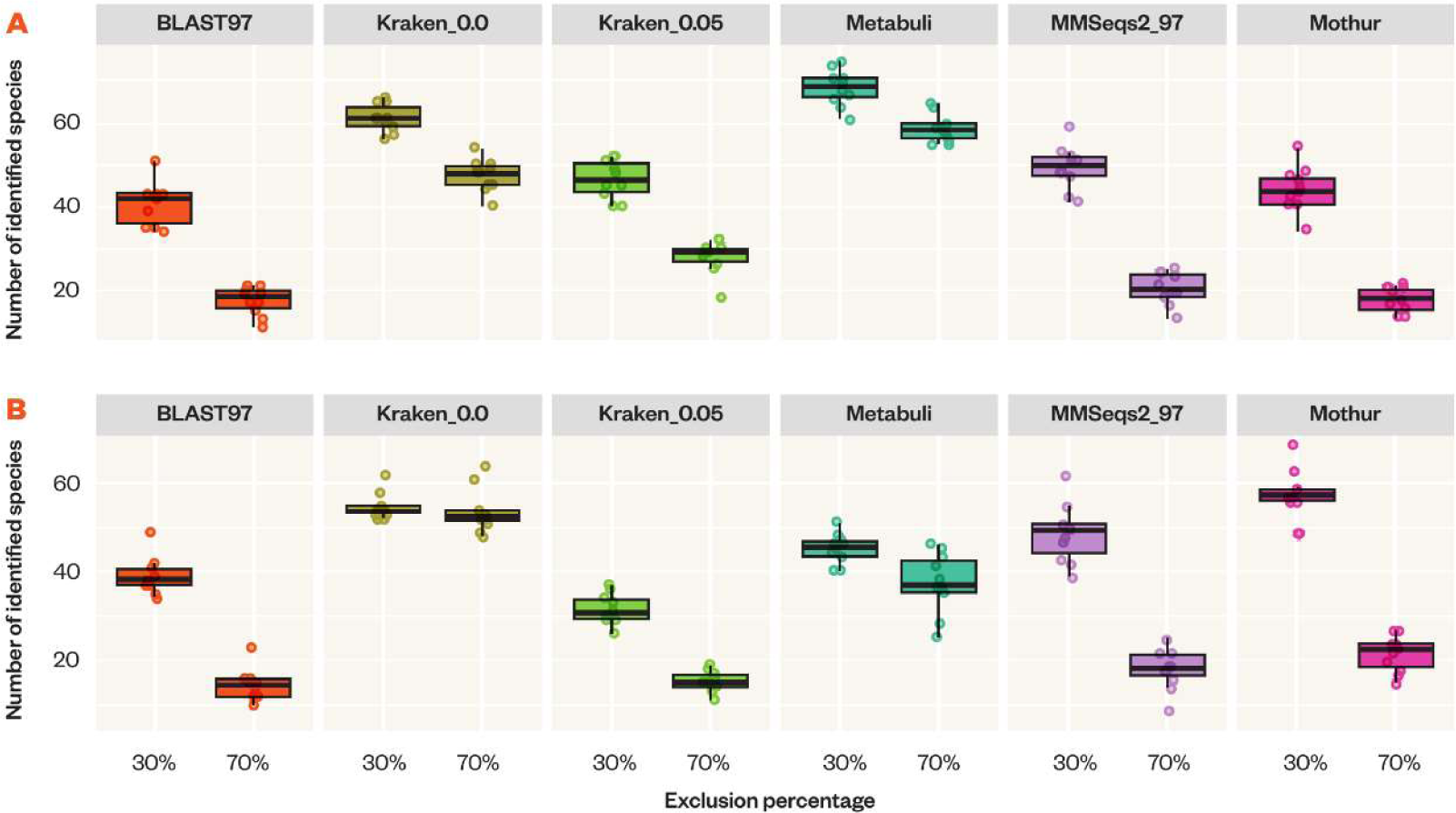
Species diversity as measured by the number of detected species across five classifiers for 30% and 70% family-level exclusion databases for 12S_Miya (A) and COI_Leray (B). Lower-case letters denote statistically significant different groups as detected by Tukey’s test (Honestly Significant Distance) after correcting p-values for multiple testing.

When using exclusion databases within the negative control datasets, four classifiers (custom Naive Bayes Classifier, Qiime2, DADA2Tax, and Mothur) began to report false positive marine vertebrate hits despite the input queries being bacterial 16S genes (Table 4). When 70% of the database was excluded, Qiime2 and CustomNBC mis-identified bacterial species for an average of 13 and 9 marine vertebrate ASVs respectively. When 50% of families were excluded, 50% of the species labels for ASVs detected by CustomNBC were false positives. When 30% of the database was excluded, only CustomNBC mis-identified marine species for 29% of bacterial ASVs. Across all percentages for COI_Leray, DADA2Tax misidentified only one species. No other classifier detected any marine vertebrate species in the bacterial Greengenes or the random database.

**Table 4:** Comparison of false positive rates among all classifiers that detected false positive species in the negative bacterial test datasets (Naive Bayes Classifier (CustomNBC), Qiime2, Mothur, and DADA2Tax) across exclusion databases (30%, 50%, and 70%). The lowest, highest, and median number of false positive hits across ten randomly excluded subsets of families is shown. Classifier/exclusion databases with no false positive hits are not shown. As we used 100 simulated queries from Greengenes the count is also the percentage.

| Gene | Classifier | Percentage of families randomly excluded | Lowest false positive hits | Highest false positive hits | Median false positive hits |
| --- | --- | --- | --- | --- | --- |
| 16S_Berry | CustomNBC | 30% | 4 | 53 | 28.5 |
|  |  | 50% | 30 | 80 | 44.5 |
|  |  | 70% | 8 | 10 | 9 |
|  | Qiime2 | 70% | 4 | 22 | 13 |
| COI_Leray | CustomNBC | 30% | 1 | 34 | 6 |
|  |  | 50% | 2 | 16 | 9 |
|  |  | 70% | 1 | 82 | 8 |
|  | DADA2Tax | 30% | 1 | 1 | 1 |
|  |  | 50% | 1 | 1 | 1 |
|  |  | 70% | 1 | 1 | 1 |
|  | Mothur | 50% | 1 | 1 | 1 |
|  |  | 70% | 2 | 2 | 2 |
|  | Qiime2 | 30% | 2 | 3 | 2.5 |
|  |  | 50% | 12 | 45 | 32 |
|  |  | 70% | 42 | 67 | 52.5 |

The false positive species detected by Qiime2 when trained on 16S and queried using Greengenes for all ASVs was *Nexilosus latifrons*. This is likely an artifact of the training process, as most implementations of Naive Bayes Classifiers start with random low probabilities for all potential species labels. It may be that in this case, *N. latifrons* was randomly assigned the highest class probability. CustomNBC, DADA2Tax, and Mothur classifier additionally found different false positive species, including *Sillago sihama, Tursiops aduncus, Monopterus albus*, *Mobula rochebrunei*, and *Plestiodon brevirostris*, which is also likely an outcome of the random label probability initialisation. There were no false positives detected by the classifiers using the 12S_Miya database as reference.

## Discussion

Here, we have demonstrated that sequence-based classifiers outperform Naive Bayes-based classifiers for12S_Miya and 16S_Berry datasets, while the opposite trend was observed for COI_Leray dataset.

For 12S_Miya and 16S_Berry, MMSeqs2 and Metabuli outperformed BLAST with 10% and 11% higher F1-scores respectively, while in the COI_Leray dataset, Naive Bayes Classifiers and MMSeqs2 outperformed BLAST by 11%. To our knowledge, there are no eDNA studies employing MMSeqs2 or Metabuli for taxonomic assignment. Both Metabuli and MMSeqs2 are under active development so future improvements may see one tool outperform the other consistently.

In COI_Leray, Naive Bayes Classifiers outperformed the sequence-based classifiers but at the cost of occasionally including false positive marine vertebrates with bacterial queries in negative control tests. Naive Bayes Classifiers outperforming other classifiers in COI is in line with previous studies (Gardner et al., 2019, Mathon et al., 2021) but to our knowledge no eDNA studies have evaluated MMSeqs2 or Metabuli. We therefore recommend that researchers utilising eDNA classification tools evaluate MMSeqs2 and Metabuli in their 12S, 16S, and COI classification tasks to maximise species-level assignments while minimising false positives, while always carefully evaluating the predicted species for the likelihood to appear in the sampled area, depth, or season.

In the 12S_Miya and 16S_Berry datasets, Naive Bayes Classifier-based tools such as Qiime2 and the custom Naive Bayes Classifier performed worse in comparison to the k-mer-based classifiers, contrary to previous publications in bacteria (Gardner et al., 2019, Ziemski et al., 2021). This is likely due to lower numbers and lower diversity in training data available for marine vertebrates compared to more commonly evaluated bacteria. It may also be due to the to the difference in PCR product length. Simulated COI_Leray primer products were 256 bp long on average whereas 12S_Miya and 16S_Berry simulated primer products were only 171 bp and 203 bp long respectively. A greater number of base-pairs in COI_Leray products (∼53-85) facilitated longer alignments, which in turn resulted in more species-level assignments.

However, an interesting observation with the COI_Leray primer products is the PCR product surpassed the collective length of the forward and reverse reads. This causes a region in the middle of COI to be omitted from the denoised ASV, leading to disparities in the alignment. No sequences align using DADA2Spec because it performs full sequence alignment which fails with the missing region in the primer product. For the COI_Leray sequences, BLAST may generate two high-scoring pairs (HSPs) per ASV in COI with roughly 50% coverage whereas it reports only one HSP in 12S and 16S. Simple filtering on HSP query length percentage, often close to 100%, may accidentally remove these hits. We advise researchers working with COI_Leray and other primers generating an incomplete PCR product to always visually evaluate alignments to increase confidence in species-level assignments.

Differences in genetic diversity across the three marker genes make a true comparison of taxonomic markers hard, as it is not possible to separate the impact of genetic diversity for a specific marker gene and the impact of the taxonomic classifier implementation choices on the final taxonomic classification outcome. A truly fair taxonomic classification comparison would use simulated marker genes across a gradient of genetic diversity. Such simulations will be the focus of future studies, as we first need to evaluate the marker genes currently being deployed in eDNA-based biodiversity studies.

The bacterial false positive controls reveal limitations in hyperparameter optimisation, where increased classifier optimisation in CustomNBC resulted in poorer generalisation and higher false positive rates. This situation is indicative of overfitting, where a classifier lacks sufficient training examples leading to confident, but false outcomes. One solution is to complete reference databases, as Naive Bayes Classifiers tend to overfit less when the reference library is more comprehensive. Similarly, the exclusion tests show that with complete reference databases, researchers can expect few or no false positive species assignments.

Incomplete reference databases are a known issue in eDNA-based studies (Somervuo et al., 2017, Marques et al., 2021, Blackman et al., 2023). Reference databases are known to be highly incomplete for fish (De Jong) hampering the completeness of eDNA-based insights. Current efforts are underway to sequence the genomes of every living species, such as the Earth BioGenome Project, the Vertebrate Genome Project, Fish10K, and Ocean Genomes, among many others.

Our random exclusion database results show that the use of negative control sets in amplicon-based studies and the significance of assessing primer pairs for specificity to their target species are needed to fully assess taxonomic classification. If these tests are not carried out, false positive findings will increase. For example, 18S-based amplicon studies focusing on marine vertebrates may accidentally sequence bacterial ASVs due to non-specific primer binding (Kumar et al., 2022). As we have shown, these bacterial ASVs may then be falsely classified as marine vertebrates when classifier hyperparameters are optimised using incomplete reference databases. These false positives can have significant implications in data interpretation and management outcomes. For instance, one of the random false-positive labels is *Tursiops aduncus*, the Indo-Pacific bottlenose dolphin, currently classified as near threatened (NT) (Braulik et al., 2019) and listed on CITES Appendix II (threatened by international trade). Consequently, these false positives and misleading labels could have erroneous influences on management decisions.

In line with previous observations in amplicon-based eDNA studies, assigning 100% of species-labels to ASVs remains unattainable due to the high sequence similarity of the candidate region between species of the same genus or family. In 12S_Miya and 16S_Berry, our study managed to assign correct species-level labels to a maximum of 62% of sequences (median MMSeqs2), only slightly surpassing the 60% species-level allocation reported in prior research (Baetscher et al., 2021). However, in COI_Leray, NBC-based classifiers could assign species-level labels for 85% of ASVs. Increasing identity cutoffs from 97% to 100% as recommended previously in bacterial studies (Edgar, 2018), slightly increased the number of species-level assignments. Therefore, we suggest a percentage identity cutoff close to 100% to enhance species-level assignments when using BLAST, but do not advocate for such high cut-offs when using MMSeqs2 as MMSeqs2_100 performed poorer than MMSeqs2_97 in almost all primer and control combinations except one (precision, 12S_Miya).

We focused only on taxonomic classification accuracy, unlike other studies that also assessed pipeline species-level read counts (Mathon et al., 2021). Tools in this space have also been shown to have an impact on biological insights (Nearing et al., 2022). We decided not to simulate different read counts as many abiotic and biotic factors influence eDNA read counts (Deiner et al., 2017, Yates et al., 2019, Rourke et al., 2022). Given that factors influencing eDNA read counts per species and marine environment have not been fully mapped, we concluded that designing a meaningful synthetic dataset for this purpose is presently not possible. Future studies can focus simulating of eDNA read levels once a comprehensive understanding of these influencing factors has been achieved.

At present, the majority of eDNA research targets mitochondrial marker genes, imposing constraints on the depth of biological insights attainable from this data. For example, many species within a genus or family possess marker genes that are almost or completely identical. As demonstrated previously, certain taxonomic classifiers tend to over-predict the number of species when marker genes are identical. Notably, fish families like Lutjanidae cannot be properly distinguished by any classifiers in the 16S dataset. One potential solution may involve employing longer sequences or even full-length genes in eDNA metabarcoding. Novel metabarcoding approaches targeting entire gene sequences improve distinction among similar species (Deiner et al., 2017, Doorenspleet et al., 2021). Future work will evaluate the impact of longer or full-length sequences on classifier performance and outcomes.

An exciting alternative to metabarcoding is shotgun metagenomics, extensively used to investigate microbial ecosystems (Rinke et al., 2013, Biller et al., 2018, Tully et al., 2018). In these studies, shotgun metagenomics is more feasible as bacterial genomes are smaller than marine vertebrate genomes. To this end, many metagenomics bioinformatics tools and pipelines focusing on microbial genomes have been released such as *Atlas* (Kieser et al., 2020) and *nf-core/mag* (Krakau et al., 2022). However, as marine vertebrate genomes are much larger and possess complex genome structures, no specialised binning tools currently exist and no studies evaluating different binning and classifier approaches in the context of marine vertebrates exist. The lack of these studies hampers the use of metagenomics methodologies in marine vertebrate eDNA studies.

Ongoing research focusing on whole-genome shotgun-sequencing approaches promises to offer more comprehensive insights into the species detected in eDNA samples, particularly when metabarcoding candidate genes fail to distinguish species or population structures. Although studies evaluating shotgun sequencing in eDNA contexts have emerged (Rinke et al., 2013, Manu and Umapathy, 2023), metabarcoding studies will remain the baseline for shotgun sequencing studies. Through our findings, we hope that the results presented here will increase confidence in metabarcoding-based ecological studies.

## Conclusion

Here we have comprehensively evaluated taxonomic classifiers using positive and negative control datasets and random database exclusions. Our findings underscore the more accurate performance of MMSeqs2 and Metabuli, evident in their consistently accurate species-level taxonomic classifications and minimized false positives, particularly in scenarios with incomplete reference databases. We therefore advocate for the widespread adoption of MMSeqs2 or Metabuli as primary tools for taxonomic classification in eDNA studies. However, further research and validation across varied environmental contexts are essential to confirm the applicability of these findings and address potential challenges to implementation.

## Acknowledgments

This work was supported by resources provided by the Pawsey Supercomputing Research Centre with funding from the Australian Government and the Government of Western Australia. We thank Associate Professor Siavash Mirarab for helpful discussion and feedback.

## Data availability statement

All code to run and parse the output of the taxonomic classifier, all code to analyse the results, all taxonomic classifier results, and simulated read datasets are available at https://github.com/MinderooFoundation/OceanOmics-classifier-comparison/. The simulation pipeline is available at https://github.com/nf-core/readsimulator. The code to curate the reference databases and the reference databases themselves are available at https://github.com/MinderooFoundation/OceanOmics-AmpliconReference.

## Benefit-Sharing statement

Benefits Generated: Benefits from this research accrue from the sharing of our data and results on public databases as described above.

## Author Contributions

PEB carried out the taxonomic profiling and evaluated profiler accuracy. AB implemented the Nextflow pipelines. PEB, PG, SC, EJR, and ASM assisted with database curation and curation steps. PM designed and drew all figures. GN, SC, MEA, AK, MWF, and MC assisted with interpretation of results and manuscript writing and editing. PEB, SB, and SR co-designed the study. All authors contributed critically to manuscript drafts and gave final approval for publication.

