## Supplementary Materials for "A comprehensive evaluation of taxonomic classifiers in marine vertebrate eDNA studies"

### Supplementary Figures

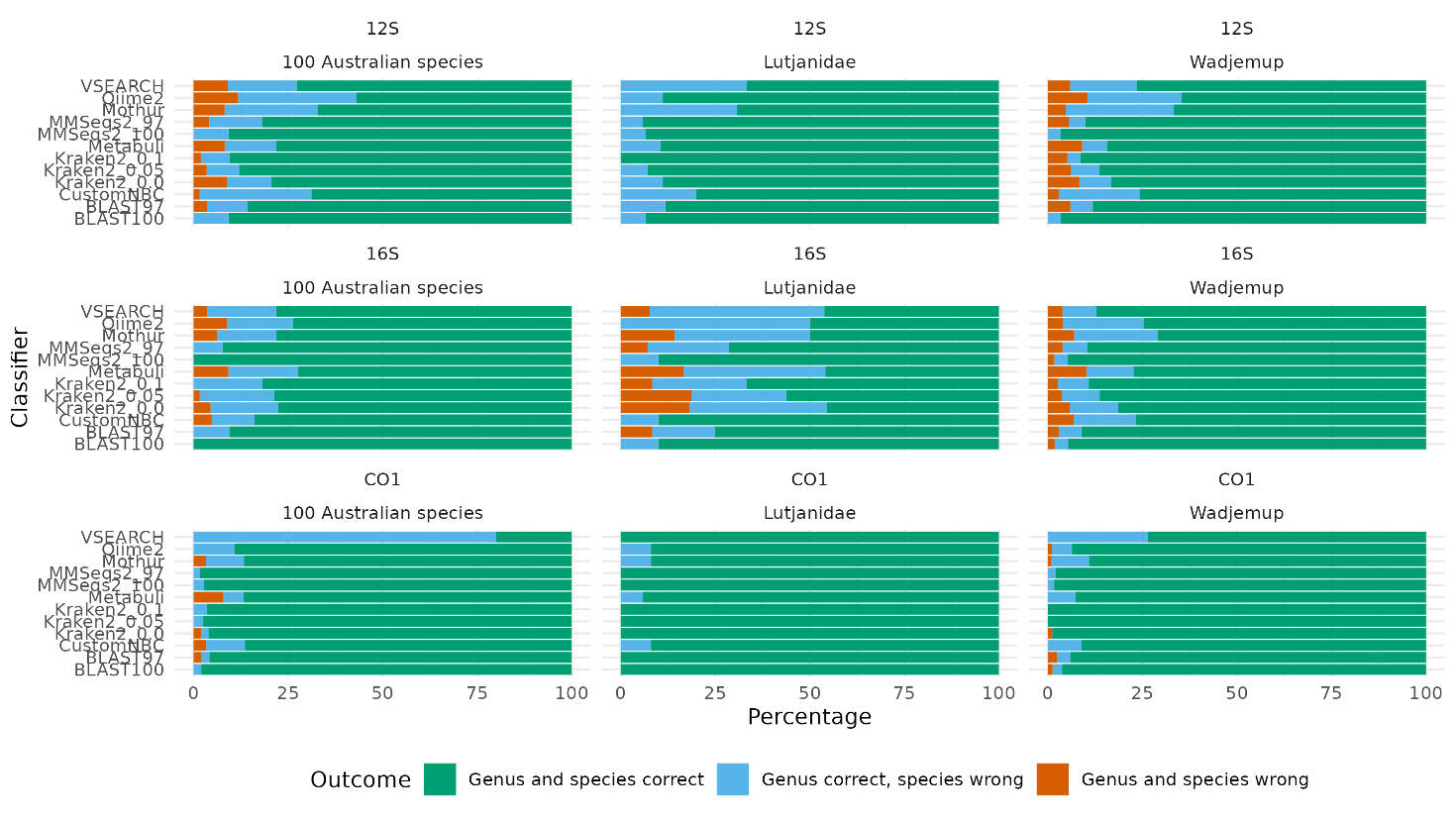

Figure S1: Error types across all databases and classifiers

### Supplementary Notes

#### Supplementary Note 1: Full Entrez search terms for 12S, 16S, and CO1

### 12S

esearch -db nuccore -query "(txid29146[ORGN] OR txid107764[ORGN] OR txid54907[ORGN] OR txid88651[ORGN] OR txid134615[ORGN] OR txid7850[ORGN] OR txid1489899[ORGN] OR txid205120[ORGN] OR txid84619[ORGN] OR txid1072510[ORGN] OR txid7934[ORGN] OR txid8279[ORGN] OR txid241819[ORGN] OR txid990930[ORGN] OR txid178227[ORGN] OR txid82890[ORGN] OR txid83881[ORGN] OR txid303732[ORGN] OR txid170194[ORGN] OR txid117867[ORGN] OR txid31017[ORGN] OR txid316123[ORGN] OR txid163127[ORGN] OR txid143304[ORGN] OR txid69128[ORGN] OR txid81361[ORGN] OR txid150446[ORGN] OR txid31024[ORGN] OR txid490307[ORGN] OR txid88658[ORGN] OR txid463601[ORGN] OR txid1489799[ORGN] OR txid172132[ORGN] OR txid1489833[ORGN] OR txid8065[ORGN] OR txid94935[ORGN] OR txid178230[ORGN] OR txid1489942[ORGN] OR txid88661[ORGN] OR txid56718[ORGN] OR txid8253[ORGN] OR txid36203[ORGN] OR txid582432[ORGN] OR txid412621[ORGN] OR txid215350[ORGN] OR txid30757[ORGN] OR txid181401[ORGN] OR txid30850[ORGN] OR txid270592[ORGN] OR txid30908[ORGN] OR txid7865[ORGN] OR txid8157[ORGN] OR txid94930[ORGN] OR txid7805[ORGN] OR txid7802[ORGN] OR txid215353[ORGN] OR txid181420[ORGN] OR txid206122[ORGN] OR txid1489924[ORGN] OR txid163122[ORGN] OR txid117924[ORGN] OR txid270595[ORGN] OR txid206106[ORGN] OR txid88666[ORGN] OR txid30828[ORGN] OR txid270610[ORGN] OR txid29142[ORGN] OR txid181415[ORGN] OR txid206140[ORGN] OR txid7869[ORGN] OR txid94304[ORGN] OR txid82793[ORGN] OR txid27583[ORGN] OR txid118146[ORGN] OR txid68511[ORGN] OR txid76914[ORGN] OR txid30943[ORGN] OR txid57768[ORGN] OR txid55118[ORGN] OR txid391217[ORGN] OR txid178225[ORGN] OR txid7941[ORGN] OR txid27766[ORGN] OR txid8092[ORGN] OR txid270598[ORGN] OR txid556237[ORGN] OR txid30947[ORGN] OR txid1545868[ORGN] OR txid94922[ORGN] OR txid117914[ORGN] OR txid30469[ORGN] OR txid118158[ORGN] OR txid412631[ORGN] OR txid390325[ORGN] OR txid31034[ORGN] OR txid88680[ORGN] OR txid215389[ORGN] OR txid75380[ORGN] OR txid365061[ORGN] OR txid173245[ORGN] OR txid7832[ORGN] OR txid86197[ORGN] OR txid7926[ORGN] OR txid181462[ORGN] OR txid43062[ORGN] OR txid215365[ORGN] OR txid75381[ORGN] OR txid223796[ORGN] OR txid117921[ORGN] OR txid630724[ORGN] OR txid55115[ORGN] OR txid172143[ORGN] OR txid76072[ORGN] OR txid206152[ORGN] OR txid1545873[ORGN] OR txid130264[ORGN] OR txid27767[ORGN] OR txid2748623[ORGN] OR txid274463[ORGN] OR txid242956[ORGN] OR txid172114[ORGN] OR txid30507[ORGN] OR txid603452[ORGN] OR txid443779[ORGN] OR txid2800422[ORGN] OR txid178224[ORGN] OR txid63826[ORGN] OR txid8220[ORGN] OR txid30764[ORGN] OR txid48439[ORGN] OR txid2025516[ORGN] OR txid31099[ORGN] OR txid86359[ORGN] OR txid30840[ORGN] OR txid143897[ORGN] OR txid376643[ORGN] OR txid88694[ORGN] OR txid40580[ORGN] OR txid7790[ORGN] OR txid7855[ORGN] OR txid117842[ORGN] OR txid242959[ORGN] OR txid47697[ORGN] OR txid30986[ORGN] OR txid390339[ORGN] OR txid117889[ORGN] OR txid760215[ORGN] OR txid172118[ORGN] OR txid1489917[ORGN] OR txid27768[ORGN] OR txid30842[ORGN] OR txid215393[ORGN] OR txid30843[ORGN] OR txid8247[ORGN] OR txid30845[ORGN] OR txid81368[ORGN] OR txid8184[ORGN] OR txid97158[ORGN] OR txid30846[ORGN] OR txid130262[ORGN] OR txid30847[ORGN] OR txid390344[ORGN] OR txid30848[ORGN] OR txid242963[ORGN] OR txid183715[ORGN] OR txid30849[ORGN] OR txid8071[ORGN] OR txid81371[ORGN] OR txid30850[ORGN] OR txid30761[ORGN] OR txid8061[ORGN] OR txid30851[ORGN] OR txid57976[ORGN] OR txid7930[ORGN] OR txid88706[ORGN] OR txid181423[ORGN] OR txid30758[ORGN] OR txid30700[ORGN] OR txid30852[ORGN] OR txid8061[ORGN] OR txid170197[ORGN] OR txid908954[ORGN] OR txid57977[ORGN] OR txid31029[ORGN] OR txid143346[ORGN] OR txid88673[ORGN] OR txid30853[ORGN] OR txid30759[ORGN] OR txid118143[ORGN] OR txid8189[ORGN] OR txid30854[ORGN] OR txid7944[ORGN] OR txid46660[ORGN] OR txid68515[ORGN] OR txid1158979[ORGN] OR txid7762[ORGN] OR txid40665[ORGN] OR txid118167[ORGN] OR txid30857[ORGN] OR txid412628[ORGN] OR txid88699[ORGN] OR txid274689[ORGN] OR txid118175[ORGN] OR txid215399[ORGN] OR txid55111[ORGN] OR txid172126[ORGN] OR txid241355[ORGN] OR txid7802[ORGN] OR txid215327[ORGN] OR txid242930[ORGN] OR txid54912[ORGN] OR txid94930[ORGN] OR txid170204[ORGN] OR txid270589[ORGN] OR txid30858[ORGN] OR txid168096[ORGN] OR txid31101[ORGN] OR txid27723[ORGN] OR txid31025[ORGN] OR txid428447[ORGN] OR txid117915[ORGN] OR txid172138[ORGN] OR txid171414[ORGN] OR txid1213740[ORGN] OR txid509847[ORGN] OR txid31102[ORGN] OR txid178226[ORGN] OR txid215337[ORGN] OR txid30859[ORGN] OR txid30860[ORGN] OR txid1545905[ORGN] OR txid8162[ORGN] OR txid178231[ORGN] OR txid390366[ORGN] OR txid210573[ORGN] OR txid206137[ORGN] OR txid30987[ORGN] OR txid170200[ORGN] OR txid117840[ORGN] OR txid30861[ORGN] OR txid8256[ORGN] OR txid30997[ORGN] OR txid490375[ORGN] OR txid81383[ORGN] OR txid30917[ORGN] OR txid112726[ORGN] OR txid30862[ORGN] OR txid30863[ORGN] OR txid30864[ORGN] OR txid30865[ORGN] OR txid55129[ORGN] OR txid260504[ORGN] OR txid55133[ORGN] OR txid30949[ORGN] OR txid1490017[ORGN] OR txid57978[ORGN] OR txid30866[ORGN] OR txid160493[ORGN] OR txid172661[ORGN] OR txid172135[ORGN] OR txid30976[ORGN] OR txid30867[ORGN] OR txid30475[ORGN] OR txid81387[ORGN] OR txid130260[ORGN] OR txid259919[ORGN] OR txid117884[ORGN] OR txid7860[ORGN] OR txid134992[ORGN] OR txid490374[ORGN] OR txid88709[ORGN] OR txid136488[ORGN] OR txid195629[ORGN] OR txid8247[ORGN] OR txid30869[ORGN] OR txid30870[ORGN] OR txid94935[ORGN] OR txid8224[ORGN] OR txid30885[ORGN] OR txid172121[ORGN] OR txid8108[ORGN] OR txid7826[ORGN] OR txid274692[ORGN] OR txid30871[ORGN] OR txid87098[ORGN] OR txid274712[ORGN] OR txid75382[ORGN] OR txid30874[ORGN] OR txid30948[ORGN] OR txid270524[ORGN] OR txid117916[ORGN] OR txid8169[ORGN] OR txid27769[ORGN] OR txid376648[ORGN] OR txid30511[ORGN] OR txid55140[ORGN] OR txid378069[ORGN] OR txid473342[ORGN] OR txid68500[ORGN] OR txid123351[ORGN] OR txid722446[ORGN] OR txid274706[ORGN] OR txid118152[ORGN] OR txid43698[ORGN] OR txid72045[ORGN] OR txid81390[ORGN] OR txid30875[ORGN] OR txid242970[ORGN] OR txid316148[ORGN] OR txid31031[ORGN] OR txid210577[ORGN] OR txid1489881[ORGN] OR txid30480[ORGN] OR txid30876[ORGN] OR txid96776[ORGN] OR txid81393[ORGN] OR txid31027[ORGN] OR txid31026[ORGN] OR txid7839[ORGN] OR txid27770[ORGN] OR txid270659[ORGN] OR txid27774[ORGN] OR txid300225[ORGN] OR txid57771[ORGN] OR txid70846[ORGN] OR txid40662[ORGN] OR txid81374[ORGN] OR txid8243[ORGN] OR txid75383[ORGN] OR txid31103[ORGN] OR txid1489918[ORGN] OR txid8193[ORGN] OR txid56259[ORGN] OR txid42164[ORGN] OR txid28683[ORGN] OR txid8830[ORGN] OR txid8899[ORGN] OR txid30558[ORGN] OR txid9765[ORGN] OR txid85103[ORGN] OR txid8907[ORGN] OR txid8465[ORGN] OR txid8493[ORGN] OR txid9726[ORGN] OR txid27792[ORGN] OR txid37067[ORGN] OR txid29135[ORGN] OR txid8602[ORGN] OR txid30443[ORGN] OR txid43314[ORGN] OR txid37576[ORGN] OR txid184169[ORGN] OR txid85545[ORGN] OR txid8910[ORGN] OR txid226940[ORGN] OR txid9702[ORGN] OR txid56260[ORGN] OR txid30444[ORGN] OR txid37076[ORGN] OR txid30445[ORGN] OR txid9206[ORGN] OR txid9709[ORGN] OR txid9740[ORGN] OR txid9750[ORGN] OR txid30448[ORGN] OR txid30451[ORGN] OR txid425629[ORGN] OR txid66056[ORGN] OR txid8917[ORGN] OR txid9231[ORGN] OR txid54049[ORGN] OR txid30446[ORGN] OR txid33574[ORGN]) AND (Small subunit [Title] OR 12S[Title] OR 12S ribosomal RNA[Title] OR 12S rRNA[Title]) NOT environmental sample[Title] NOT environmental samples[Title] NOT environmental[Title] NOT uncultured[Title] NOT unclassified[Title] NOT unidentified[Title] NOT unverified[Title] " | efetch -format fasta > 12s_fish_nuccore.fasta

esearch -db gene -query "(txid29146[ORGN] OR txid107764[ORGN] OR txid54907[ORGN] OR txid134615[ORGN] OR txid7850[ORGN] OR txid1489899[ORGN] OR txid205120[ORGN] OR txid84619[ORGN] OR txid1072510[ORGN] OR txid7934[ORGN] OR txid8279[ORGN] OR txid83881[ORGN] OR txid303732[ORGN] OR txid170194[ORGN] OR txid117867[ORGN] OR txid31017[ORGN] OR txid316123[ORGN] OR txid163127[ORGN] OR txid143304[ORGN] OR txid69128[ORGN] OR txid81361[ORGN] OR txid150446[ORGN] OR txid31024[ORGN] OR txid490307[ORGN] OR txid88658[ORGN] OR txid463601[ORGN] OR txid1489799[ORGN] OR txid8065[ORGN] OR txid94935[ORGN] OR txid88661[ORGN] OR txid56718[ORGN] OR txid8253[ORGN] OR txid36203[ORGN] OR txid215350[ORGN] OR txid30757[ORGN] OR txid181401[ORGN] OR txid30850[ORGN] OR txid270592[ORGN] OR txid30908[ORGN] OR txid7865[ORGN] OR txid8157[ORGN] OR txid94930[ORGN] OR txid7805[ORGN] OR txid7802[ORGN] OR txid181420[ORGN] OR txid206122[ORGN] OR txid163122[ORGN] OR txid270595[ORGN] OR txid206106[ORGN] OR txid88666[ORGN] OR txid30828[ORGN] OR txid270610[ORGN] OR txid29142[ORGN] OR txid181415[ORGN] OR txid206140[ORGN] OR txid7869[ORGN] OR txid94304[ORGN] OR txid27583[ORGN] OR txid118146[ORGN] OR txid68511[ORGN] OR txid76914[ORGN] OR txid30943[ORGN] OR txid55118[ORGN] OR txid391217[ORGN] OR txid7941[ORGN] OR txid27766[ORGN] OR txid8092[ORGN] OR txid556237[ORGN] OR txid30947[ORGN] OR txid94922[ORGN] OR txid117914[ORGN] OR txid30469[ORGN] OR txid118158[ORGN] OR txid412631[ORGN] OR txid31034[ORGN] OR txid88680[ORGN] OR txid215389[ORGN] OR txid75380[ORGN] OR txid365061[ORGN] OR txid173245[ORGN] OR txid86197[ORGN] OR txid7926[ORGN] OR txid181462[ORGN] OR txid43062[ORGN] OR txid215365[ORGN] OR txid75381[ORGN] OR txid223796[ORGN] OR txid117921[ORGN] OR txid55115[ORGN] OR txid76072[ORGN] OR txid206152[ORGN] OR txid130264[ORGN] OR txid27767[ORGN] OR txid2748623[ORGN] OR txid274463[ORGN] OR txid242956[ORGN] OR txid30507[ORGN] OR txid603452[ORGN] OR txid63826[ORGN] OR txid8220[ORGN] OR txid30764[ORGN] OR txid48439[ORGN] OR txid86359[ORGN] OR txid30840[ORGN] OR txid143897[ORGN] OR txid376643[ORGN] OR txid88694[ORGN] OR txid40580[ORGN] OR txid7790[ORGN] OR txid7855[ORGN] OR txid242959[ORGN] OR txid47697[ORGN] OR txid30986[ORGN] OR txid390339[ORGN] OR txid1489917[ORGN] OR txid27768[ORGN] OR txid30842[ORGN] OR txid215393[ORGN] OR txid30843[ORGN] OR txid8247[ORGN] OR txid30845[ORGN] OR txid81368[ORGN] OR txid8184[ORGN] OR txid30846[ORGN] OR txid30848[ORGN] OR txid242963[ORGN] OR txid183715[ORGN] OR txid30849[ORGN] OR txid8071[ORGN] OR txid30850[ORGN] OR txid30761[ORGN] OR txid8061[ORGN] OR txid30851[ORGN] OR txid57976[ORGN] OR txid7930[ORGN] OR txid88706[ORGN] OR txid181423[ORGN] OR txid30700[ORGN] OR txid30852[ORGN] OR txid8061[ORGN] OR txid170197[ORGN] OR txid57977[ORGN] OR txid31029[ORGN] OR txid143346[ORGN] OR txid88673[ORGN] OR txid30853[ORGN] OR txid30759[ORGN] OR txid118143[ORGN] OR txid8189[ORGN] OR txid30854[ORGN] OR txid7944[ORGN] OR txid46660[ORGN] OR txid68515[ORGN] OR txid1158979[ORGN] OR txid7762[ORGN] OR txid40665[ORGN] OR txid118167[ORGN] OR txid30857[ORGN] OR txid412628[ORGN] OR txid88699[ORGN] OR txid118175[ORGN] OR txid215399[ORGN] OR txid55111[ORGN] OR txid241355[ORGN] OR txid7802[ORGN] OR txid215327[ORGN] OR txid242930[ORGN] OR txid54912[ORGN] OR txid94930[ORGN] OR txid170204[ORGN] OR txid270589[ORGN] OR txid30858[ORGN] OR txid168096[ORGN] OR txid31101[ORGN] OR txid27723[ORGN] OR txid31025[ORGN] OR txid172138[ORGN] OR txid171414[ORGN] OR txid31102[ORGN] OR txid215337[ORGN] OR txid30859[ORGN] OR txid30860[ORGN] OR txid1545905[ORGN] OR txid8162[ORGN] OR txid178231[ORGN] OR txid30987[ORGN] OR txid170200[ORGN] OR txid117840[ORGN] OR txid30861[ORGN] OR txid8256[ORGN] OR txid30997[ORGN] OR txid81383[ORGN] OR txid30917[ORGN] OR txid30862[ORGN] OR txid30863[ORGN] OR txid30864[ORGN] OR txid30865[ORGN] OR txid55129[ORGN] OR txid260504[ORGN] OR txid55133[ORGN] OR txid30949[ORGN] OR txid57978[ORGN] OR txid30866[ORGN] OR txid172661[ORGN] OR txid30976[ORGN] OR txid30867[ORGN] OR txid30475[ORGN] OR txid130260[ORGN] OR txid259919[ORGN] OR txid117884[ORGN] OR txid7860[ORGN] OR txid134992[ORGN] OR txid490374[ORGN] OR txid88709[ORGN] OR txid136488[ORGN] OR txid195629[ORGN] OR txid8247[ORGN] OR txid30869[ORGN] OR txid30870[ORGN] OR txid94935[ORGN] OR txid8224[ORGN] OR txid8108[ORGN] OR txid7826[ORGN] OR txid274692[ORGN] OR txid30871[ORGN] OR txid87098[ORGN] OR txid75382[ORGN] OR txid30874[ORGN] OR txid30948[ORGN] OR txid270524[ORGN] OR txid117916[ORGN] OR txid8169[ORGN] OR txid27769[ORGN] OR txid376648[ORGN] OR txid30511[ORGN] OR txid55140[ORGN] OR txid378069[ORGN] OR txid68500[ORGN] OR txid123351[ORGN] OR txid274706[ORGN] OR txid118152[ORGN] OR txid43698[ORGN] OR txid72045[ORGN] OR txid81390[ORGN] OR txid30875[ORGN] OR txid242970[ORGN] OR txid316148[ORGN] OR txid31031[ORGN] OR txid210577[ORGN] OR txid30480[ORGN] OR txid30876[ORGN] OR txid96776[ORGN] OR txid81393[ORGN] OR txid31027[ORGN] OR txid31026[ORGN] OR txid7839[ORGN] OR txid27770[ORGN] OR txid270659[ORGN] OR txid27774[ORGN] OR txid300225[ORGN] OR txid70846[ORGN] OR txid8243[ORGN] OR txid75383[ORGN] OR txid31103[ORGN] OR txid1489918[ORGN] OR txid8193[ORGN] OR txid56259[ORGN] OR txid42164[ORGN] OR txid28683[ORGN] OR txid8830[ORGN] OR txid8899[ORGN] OR txid30558[ORGN] OR txid9765[ORGN] OR txid85103[ORGN] OR txid8907[ORGN] OR txid8465[ORGN] OR txid8493[ORGN] OR txid9726[ORGN] OR txid27792[ORGN] OR txid37067[ORGN] OR txid29135[ORGN] OR txid8602[ORGN] OR txid30443[ORGN] OR txid37576[ORGN] OR txid184169[ORGN] OR txid85545[ORGN] OR txid8910[ORGN] OR txid226940[ORGN] OR txid9702[ORGN] OR txid56260[ORGN] OR txid30444[ORGN] OR txid37076[ORGN] OR txid30445[ORGN] OR txid9206[ORGN] OR txid9709[ORGN] OR txid9740[ORGN] OR txid9750[ORGN] OR txid30448[ORGN] OR txid30451[ORGN] OR txid425629[ORGN] OR txid66056[ORGN] OR txid8917[ORGN] OR txid9231[ORGN] OR txid54049[ORGN] OR txid30446[ORGN] OR txid33574[ORGN]) AND (Small subunit [Title] OR 12S[Title])" | efetch -format docsum |	xtract -pattern GenomicInfoType -element ChrAccVer ChrStart ChrStop | while IFS=$'\t' read acn str stp; do	efetch -db nuccore -format fasta -id "$acn" -chr_start "$str" -chr_stop "$stp"; done | sed "s/>/>12S_/" > 12s_fish_gene.fasta

### 16S

esearch -db nuccore -query "(txid29146[ORGN] OR txid107764[ORGN] OR txid54907[ORGN] OR txid88651[ORGN] OR txid134615[ORGN] OR txid7850[ORGN] OR txid1489899[ORGN] OR txid205120[ORGN] OR txid84619[ORGN] OR txid1072510[ORGN] OR txid7934[ORGN] OR txid8279[ORGN] OR txid241819[ORGN] OR txid990930[ORGN] OR txid178227[ORGN] OR txid82890[ORGN] OR txid83881[ORGN] OR txid303732[ORGN] OR txid170194[ORGN] OR txid117867[ORGN] OR txid31017[ORGN] OR txid316123[ORGN] OR txid163127[ORGN] OR txid143304[ORGN] OR txid69128[ORGN] OR txid81361[ORGN] OR txid150446[ORGN] OR txid31024[ORGN] OR txid490307[ORGN] OR txid88658[ORGN] OR txid463601[ORGN] OR txid1489799[ORGN] OR txid172132[ORGN] OR txid1489833[ORGN] OR txid8065[ORGN] OR txid94935[ORGN] OR txid178230[ORGN] OR txid1489942[ORGN] OR txid88661[ORGN] OR txid56718[ORGN] OR txid8253[ORGN] OR txid36203[ORGN] OR txid582432[ORGN] OR txid412621[ORGN] OR txid215350[ORGN] OR txid30757[ORGN] OR txid181401[ORGN] OR txid30850[ORGN] OR txid270592[ORGN] OR txid30908[ORGN] OR txid7865[ORGN] OR txid8157[ORGN] OR txid94930[ORGN] OR txid7805[ORGN] OR txid7802[ORGN] OR txid215353[ORGN] OR txid181420[ORGN] OR txid206122[ORGN] OR txid1489924[ORGN] OR txid163122[ORGN] OR txid117924[ORGN] OR txid270595[ORGN] OR txid206106[ORGN] OR txid88666[ORGN] OR txid30828[ORGN] OR txid270610[ORGN] OR txid29142[ORGN] OR txid181415[ORGN] OR txid206140[ORGN] OR txid7869[ORGN] OR txid94304[ORGN] OR txid82793[ORGN] OR txid27583[ORGN] OR txid118146[ORGN] OR txid68511[ORGN] OR txid76914[ORGN] OR txid30943[ORGN] OR txid57768[ORGN] OR txid55118[ORGN] OR txid391217[ORGN] OR txid178225[ORGN] OR txid7941[ORGN] OR txid27766[ORGN] OR txid8092[ORGN] OR txid270598[ORGN] OR txid556237[ORGN] OR txid30947[ORGN] OR txid1545868[ORGN] OR txid94922[ORGN] OR txid117914[ORGN] OR txid30469[ORGN] OR txid118158[ORGN] OR txid412631[ORGN] OR txid390325[ORGN] OR txid31034[ORGN] OR txid88680[ORGN] OR txid215389[ORGN] OR txid75380[ORGN] OR txid365061[ORGN] OR txid173245[ORGN] OR txid7832[ORGN] OR txid86197[ORGN] OR txid7926[ORGN] OR txid181462[ORGN] OR txid43062[ORGN] OR txid215365[ORGN] OR txid75381[ORGN] OR txid223796[ORGN] OR txid117921[ORGN] OR txid630724[ORGN] OR txid55115[ORGN] OR txid172143[ORGN] OR txid76072[ORGN] OR txid206152[ORGN] OR txid1545873[ORGN] OR txid130264[ORGN] OR txid27767[ORGN] OR txid2748623[ORGN] OR txid274463[ORGN] OR txid242956[ORGN] OR txid172114[ORGN] OR txid30507[ORGN] OR txid603452[ORGN] OR txid443779[ORGN] OR txid2800422[ORGN] OR txid178224[ORGN] OR txid63826[ORGN] OR txid8220[ORGN] OR txid30764[ORGN] OR txid48439[ORGN] OR txid2025516[ORGN] OR txid31099[ORGN] OR txid86359[ORGN] OR txid30840[ORGN] OR txid143897[ORGN] OR txid376643[ORGN] OR txid88694[ORGN] OR txid40580[ORGN] OR txid7790[ORGN] OR txid7855[ORGN] OR txid117842[ORGN] OR txid242959[ORGN] OR txid47697[ORGN] OR txid30986[ORGN] OR txid390339[ORGN] OR txid117889[ORGN] OR txid760215[ORGN] OR txid172118[ORGN] OR txid1489917[ORGN] OR txid27768[ORGN] OR txid30842[ORGN] OR txid215393[ORGN] OR txid30843[ORGN] OR txid8247[ORGN] OR txid30845[ORGN] OR txid81368[ORGN] OR txid8184[ORGN] OR txid97158[ORGN] OR txid30846[ORGN] OR txid130262[ORGN] OR txid30847[ORGN] OR txid390344[ORGN] OR txid30848[ORGN] OR txid242963[ORGN] OR txid183715[ORGN] OR txid30849[ORGN] OR txid8071[ORGN] OR txid81371[ORGN] OR txid30850[ORGN] OR txid30761[ORGN] OR txid8061[ORGN] OR txid30851[ORGN] OR txid57976[ORGN] OR txid7930[ORGN] OR txid88706[ORGN] OR txid181423[ORGN] OR txid30758[ORGN] OR txid30700[ORGN] OR txid30852[ORGN] OR txid8061[ORGN] OR txid170197[ORGN] OR txid908954[ORGN] OR txid57977[ORGN] OR txid31029[ORGN] OR txid143346[ORGN] OR txid88673[ORGN] OR txid30853[ORGN] OR txid30759[ORGN] OR txid118143[ORGN] OR txid8189[ORGN] OR txid30854[ORGN] OR txid7944[ORGN] OR txid46660[ORGN] OR txid68515[ORGN] OR txid1158979[ORGN] OR txid7762[ORGN] OR txid40665[ORGN] OR txid118167[ORGN] OR txid30857[ORGN] OR txid412628[ORGN] OR txid88699[ORGN] OR txid274689[ORGN] OR txid118175[ORGN] OR txid215399[ORGN] OR txid55111[ORGN] OR txid172126[ORGN] OR txid241355[ORGN] OR txid7802[ORGN] OR txid215327[ORGN] OR txid242930[ORGN] OR txid54912[ORGN] OR txid94930[ORGN] OR txid170204[ORGN] OR txid270589[ORGN] OR txid30858[ORGN] OR txid168096[ORGN] OR txid31101[ORGN] OR txid27723[ORGN] OR txid31025[ORGN] OR txid428447[ORGN] OR txid117915[ORGN] OR txid172138[ORGN] OR txid171414[ORGN] OR txid1213740[ORGN] OR txid509847[ORGN] OR txid31102[ORGN] OR txid178226[ORGN] OR txid215337[ORGN] OR txid30859[ORGN] OR txid30860[ORGN] OR txid1545905[ORGN] OR txid8162[ORGN] OR txid178231[ORGN] OR txid390366[ORGN] OR txid210573[ORGN] OR txid206137[ORGN] OR txid30987[ORGN] OR txid170200[ORGN] OR txid117840[ORGN] OR txid30861[ORGN] OR txid8256[ORGN] OR txid30997[ORGN] OR txid490375[ORGN] OR txid81383[ORGN] OR txid30917[ORGN] OR txid112726[ORGN] OR txid30862[ORGN] OR txid30863[ORGN] OR txid30864[ORGN] OR txid30865[ORGN] OR txid55129[ORGN] OR txid260504[ORGN] OR txid55133[ORGN] OR txid30949[ORGN] OR txid1490017[ORGN] OR txid57978[ORGN] OR txid30866[ORGN] OR txid160493[ORGN] OR txid172661[ORGN] OR txid172135[ORGN] OR txid30976[ORGN] OR txid30867[ORGN] OR txid30475[ORGN] OR txid81387[ORGN] OR txid130260[ORGN] OR txid259919[ORGN] OR txid117884[ORGN] OR txid7860[ORGN] OR txid134992[ORGN] OR txid490374[ORGN] OR txid88709[ORGN] OR txid136488[ORGN] OR txid195629[ORGN] OR txid8247[ORGN] OR txid30869[ORGN] OR txid30870[ORGN] OR txid94935[ORGN] OR txid8224[ORGN] OR txid30885[ORGN] OR txid172121[ORGN] OR txid8108[ORGN] OR txid7826[ORGN] OR txid274692[ORGN] OR txid30871[ORGN] OR txid87098[ORGN] OR txid274712[ORGN] OR txid75382[ORGN] OR txid30874[ORGN] OR txid30948[ORGN] OR txid270524[ORGN] OR txid117916[ORGN] OR txid8169[ORGN] OR txid27769[ORGN] OR txid376648[ORGN] OR txid30511[ORGN] OR txid55140[ORGN] OR txid378069[ORGN] OR txid473342[ORGN] OR txid68500[ORGN] OR txid123351[ORGN] OR txid722446[ORGN] OR txid274706[ORGN] OR txid118152[ORGN] OR txid43698[ORGN] OR txid72045[ORGN] OR txid81390[ORGN] OR txid30875[ORGN] OR txid242970[ORGN] OR txid316148[ORGN] OR txid31031[ORGN] OR txid210577[ORGN] OR txid1489881[ORGN] OR txid30480[ORGN] OR txid30876[ORGN] OR txid96776[ORGN] OR txid81393[ORGN] OR txid31027[ORGN] OR txid31026[ORGN] OR txid7839[ORGN] OR txid27770[ORGN] OR txid270659[ORGN] OR txid27774[ORGN] OR txid300225[ORGN] OR txid57771[ORGN] OR txid70846[ORGN] OR txid40662[ORGN] OR txid81374[ORGN] OR txid8243[ORGN] OR txid75383[ORGN] OR txid31103[ORGN] OR txid1489918[ORGN] OR txid8193[ORGN] OR txid56259[ORGN] OR txid42164[ORGN] OR txid28683[ORGN] OR txid8830[ORGN] OR txid8899[ORGN] OR txid30558[ORGN] OR txid9765[ORGN] OR txid85103[ORGN] OR txid8907[ORGN] OR txid8465[ORGN] OR txid8493[ORGN] OR txid9726[ORGN] OR txid27792[ORGN] OR txid37067[ORGN] OR txid29135[ORGN] OR txid8602[ORGN] OR txid30443[ORGN] OR txid43314[ORGN] OR txid37576[ORGN] OR txid184169[ORGN] OR txid85545[ORGN] OR txid8910[ORGN] OR txid226940[ORGN] OR txid9702[ORGN] OR txid56260[ORGN] OR txid30444[ORGN] OR txid37076[ORGN] OR txid30445[ORGN] OR txid9206[ORGN] OR txid9709[ORGN] OR txid9740[ORGN] OR txid9750[ORGN] OR txid30448[ORGN] OR txid30451[ORGN] OR txid425629[ORGN] OR txid66056[ORGN] OR txid8917[ORGN] OR txid9231[ORGN] OR txid54049[ORGN] OR txid30446[ORGN] OR txid33574[ORGN]) AND (Small subunit [Title] OR 16S[Title] OR 16S ribosomal RNA[Title] OR 16S rRNA[Title]) AND (mitochondrion[Filter] OR plastid[Filter]) NOT environmental sample[Title] NOT environmental samples[Title] NOT environmental[Title] NOT uncultured[Title] NOT unclassified[Title] NOT unidentified[Title] NOT unverified[Title] " | efetch -format fasta > 16S_fish_nuccore.fasta

esearch -db gene -query "(txid29146[ORGN] OR txid107764[ORGN] OR txid54907[ORGN] OR txid134615[ORGN] OR txid7850[ORGN] OR txid1489899[ORGN] OR txid205120[ORGN] OR txid84619[ORGN] OR txid1072510[ORGN] OR txid7934[ORGN] OR txid8279[ORGN] OR txid83881[ORGN] OR txid303732[ORGN] OR txid170194[ORGN] OR txid117867[ORGN] OR txid31017[ORGN] OR txid316123[ORGN] OR txid163127[ORGN] OR txid143304[ORGN] OR txid69128[ORGN] OR txid81361[ORGN] OR txid150446[ORGN] OR txid31024[ORGN] OR txid490307[ORGN] OR txid88658[ORGN] OR txid463601[ORGN] OR txid1489799[ORGN] OR txid8065[ORGN] OR txid94935[ORGN] OR txid88661[ORGN] OR txid56718[ORGN] OR txid8253[ORGN] OR txid36203[ORGN] OR txid215350[ORGN] OR txid30757[ORGN] OR txid181401[ORGN] OR txid30850[ORGN] OR txid270592[ORGN] OR txid30908[ORGN] OR txid7865[ORGN] OR txid8157[ORGN] OR txid94930[ORGN] OR txid7805[ORGN] OR txid7802[ORGN] OR txid181420[ORGN] OR txid206122[ORGN] OR txid163122[ORGN] OR txid270595[ORGN] OR txid206106[ORGN] OR txid88666[ORGN] OR txid30828[ORGN] OR txid270610[ORGN] OR txid29142[ORGN] OR txid181415[ORGN] OR txid206140[ORGN] OR txid7869[ORGN] OR txid94304[ORGN] OR txid27583[ORGN] OR txid118146[ORGN] OR txid68511[ORGN] OR txid76914[ORGN] OR txid30943[ORGN] OR txid55118[ORGN] OR txid391217[ORGN] OR txid7941[ORGN] OR txid27766[ORGN] OR txid8092[ORGN] OR txid556237[ORGN] OR txid30947[ORGN] OR txid94922[ORGN] OR txid117914[ORGN] OR txid30469[ORGN] OR txid118158[ORGN] OR txid412631[ORGN] OR txid31034[ORGN] OR txid88680[ORGN] OR txid215389[ORGN] OR txid75380[ORGN] OR txid365061[ORGN] OR txid173245[ORGN] OR txid86197[ORGN] OR txid7926[ORGN] OR txid181462[ORGN] OR txid43062[ORGN] OR txid215365[ORGN] OR txid75381[ORGN] OR txid223796[ORGN] OR txid117921[ORGN] OR txid55115[ORGN] OR txid76072[ORGN] OR txid206152[ORGN] OR txid130264[ORGN] OR txid27767[ORGN] OR txid2748623[ORGN] OR txid274463[ORGN] OR txid242956[ORGN] OR txid30507[ORGN] OR txid603452[ORGN] OR txid63826[ORGN] OR txid8220[ORGN] OR txid30764[ORGN] OR txid48439[ORGN] OR txid86359[ORGN] OR txid30840[ORGN] OR txid143897[ORGN] OR txid376643[ORGN] OR txid88694[ORGN] OR txid40580[ORGN] OR txid7790[ORGN] OR txid7855[ORGN] OR txid242959[ORGN] OR txid47697[ORGN] OR txid30986[ORGN] OR txid390339[ORGN] OR txid1489917[ORGN] OR txid27768[ORGN] OR txid30842[ORGN] OR txid215393[ORGN] OR txid30843[ORGN] OR txid8247[ORGN] OR txid30845[ORGN] OR txid81368[ORGN] OR txid8184[ORGN] OR txid30846[ORGN] OR txid30848[ORGN] OR txid242963[ORGN] OR txid183715[ORGN] OR txid30849[ORGN] OR txid8071[ORGN] OR txid30850[ORGN] OR txid30761[ORGN] OR txid8061[ORGN] OR txid30851[ORGN] OR txid57976[ORGN] OR txid7930[ORGN] OR txid88706[ORGN] OR txid181423[ORGN] OR txid30700[ORGN] OR txid30852[ORGN] OR txid8061[ORGN] OR txid170197[ORGN] OR txid57977[ORGN] OR txid31029[ORGN] OR txid143346[ORGN] OR txid88673[ORGN] OR txid30853[ORGN] OR txid30759[ORGN] OR txid118143[ORGN] OR txid8189[ORGN] OR txid30854[ORGN] OR txid7944[ORGN] OR txid46660[ORGN] OR txid68515[ORGN] OR txid1158979[ORGN] OR txid7762[ORGN] OR txid40665[ORGN] OR txid118167[ORGN] OR txid30857[ORGN] OR txid412628[ORGN] OR txid88699[ORGN] OR txid118175[ORGN] OR txid215399[ORGN] OR txid55111[ORGN] OR txid241355[ORGN] OR txid7802[ORGN] OR txid215327[ORGN] OR txid242930[ORGN] OR txid54912[ORGN] OR txid94930[ORGN] OR txid170204[ORGN] OR txid270589[ORGN] OR txid30858[ORGN] OR txid168096[ORGN] OR txid31101[ORGN] OR txid27723[ORGN] OR txid31025[ORGN] OR txid172138[ORGN] OR txid171414[ORGN] OR txid31102[ORGN] OR txid215337[ORGN] OR txid30859[ORGN] OR txid30860[ORGN] OR txid1545905[ORGN] OR txid8162[ORGN] OR txid178231[ORGN] OR txid30987[ORGN] OR txid170200[ORGN] OR txid117840[ORGN] OR txid30861[ORGN] OR txid8256[ORGN] OR txid30997[ORGN] OR txid81383[ORGN] OR txid30917[ORGN] OR txid30862[ORGN] OR txid30863[ORGN] OR txid30864[ORGN] OR txid30865[ORGN] OR txid55129[ORGN] OR txid260504[ORGN] OR txid55133[ORGN] OR txid30949[ORGN] OR txid57978[ORGN] OR txid30866[ORGN] OR txid172661[ORGN] OR txid30976[ORGN] OR txid30867[ORGN] OR txid30475[ORGN] OR txid130260[ORGN] OR txid259919[ORGN] OR txid117884[ORGN] OR txid7860[ORGN] OR txid134992[ORGN] OR txid490374[ORGN] OR txid88709[ORGN] OR txid136488[ORGN] OR txid195629[ORGN] OR txid8247[ORGN] OR txid30869[ORGN] OR txid30870[ORGN] OR txid94935[ORGN] OR txid8224[ORGN] OR txid8108[ORGN] OR txid7826[ORGN] OR txid274692[ORGN] OR txid30871[ORGN] OR txid87098[ORGN] OR txid75382[ORGN] OR txid30874[ORGN] OR txid30948[ORGN] OR txid270524[ORGN] OR txid117916[ORGN] OR txid8169[ORGN] OR txid27769[ORGN] OR txid376648[ORGN] OR txid30511[ORGN] OR txid55140[ORGN] OR txid378069[ORGN] OR txid68500[ORGN] OR txid123351[ORGN] OR txid274706[ORGN] OR txid118152[ORGN] OR txid43698[ORGN] OR txid72045[ORGN] OR txid81390[ORGN] OR txid30875[ORGN] OR txid242970[ORGN] OR txid316148[ORGN] OR txid31031[ORGN] OR txid210577[ORGN] OR txid30480[ORGN] OR txid30876[ORGN] OR txid96776[ORGN] OR txid81393[ORGN] OR txid31027[ORGN] OR txid31026[ORGN] OR txid7839[ORGN] OR txid27770[ORGN] OR txid270659[ORGN] OR txid27774[ORGN] OR txid300225[ORGN] OR txid70846[ORGN] OR txid8243[ORGN] OR txid75383[ORGN] OR txid31103[ORGN] OR txid1489918[ORGN] OR txid8193[ORGN] OR txid56259[ORGN] OR txid42164[ORGN] OR txid28683[ORGN] OR txid8830[ORGN] OR txid8899[ORGN] OR txid30558[ORGN] OR txid9765[ORGN] OR txid85103[ORGN] OR txid8907[ORGN] OR txid8465[ORGN] OR txid8493[ORGN] OR txid9726[ORGN] OR txid27792[ORGN] OR txid37067[ORGN] OR txid29135[ORGN] OR txid8602[ORGN] OR txid30443[ORGN] OR txid37576[ORGN] OR txid184169[ORGN] OR txid85545[ORGN] OR txid8910[ORGN] OR txid226940[ORGN] OR txid9702[ORGN] OR txid56260[ORGN] OR txid30444[ORGN] OR txid37076[ORGN] OR txid30445[ORGN] OR txid9206[ORGN] OR txid9709[ORGN] OR txid9740[ORGN] OR txid9750[ORGN] OR txid30448[ORGN] OR txid30451[ORGN] OR txid425629[ORGN] OR txid66056[ORGN] OR txid8917[ORGN] OR txid9231[ORGN] OR txid54049[ORGN] OR txid30446[ORGN] OR txid33574[ORGN]) AND (Small subunit [Title] OR 16S[Title])" | efetch -format docsum | xtract -pattern GenomicInfoType -element ChrAccVer ChrStart ChrStop | while IFS=$'\t' read acn str stp; do efetch -db nuccore -format fasta -id "$acn" -chr_start "$str" -chr_stop "$stp"; done | sed "s/>/>16S_/" > 16S_fish_gene.fasta

### CO1

esearch -db nuccore -query '(txid29146[ORGN] OR txid107764[ORGN] OR txid54907[ORGN] OR txid88651[ORGN] OR txid134615[ORGN] OR txid7850[ORGN] OR txid1489899[ORGN] OR txid205120[ORGN] OR txid84619[ORGN] OR txid1072510[ORGN] OR txid7934[ORGN] OR txid8279[ORGN] OR txid241819[ORGN] OR txid990930[ORGN] OR txid178227[ORGN] OR txid82890[ORGN] OR txid83881[ORGN] OR txid303732[ORGN] OR txid170194[ORGN] OR txid117867[ORGN] OR txid31017[ORGN] OR txid316123[ORGN] OR txid163127[ORGN] OR txid143304[ORGN] OR txid69128[ORGN] OR txid81361[ORGN] OR txid150446[ORGN] OR txid31024[ORGN] OR txid490307[ORGN] OR txid88658[ORGN] OR txid463601[ORGN] OR txid1489799[ORGN] OR txid172132[ORGN] OR txid1489833[ORGN] OR txid8065[ORGN] OR txid94935[ORGN] OR txid178230[ORGN] OR txid1489942[ORGN] OR txid88661[ORGN] OR txid56718[ORGN] OR txid8253[ORGN] OR txid36203[ORGN] OR txid582432[ORGN] OR txid412621[ORGN] OR txid215350[ORGN] OR txid30757[ORGN] OR txid181401[ORGN] OR txid30850[ORGN] OR txid270592[ORGN] OR txid30908[ORGN] OR txid7865[ORGN] OR txid8157[ORGN] OR txid94930[ORGN] OR txid7805[ORGN] OR txid7802[ORGN] OR txid215353[ORGN] OR txid181420[ORGN] OR txid206122[ORGN] OR txid1489924[ORGN] OR txid163122[ORGN] OR txid117924[ORGN] OR txid270595[ORGN] OR txid206106[ORGN] OR txid88666[ORGN] OR txid30828[ORGN] OR txid270610[ORGN] OR txid29142[ORGN] OR txid181415[ORGN] OR txid206140[ORGN] OR txid7869[ORGN] OR txid94304[ORGN] OR txid82793[ORGN] OR txid27583[ORGN] OR txid118146[ORGN] OR txid68511[ORGN] OR txid76914[ORGN] OR txid30943[ORGN] OR txid57768[ORGN] OR txid55118[ORGN] OR txid391217[ORGN] OR txid178225[ORGN] OR txid7941[ORGN] OR txid27766[ORGN] OR txid8092[ORGN] OR txid270598[ORGN] OR txid556237[ORGN] OR txid30947[ORGN] OR txid1545868[ORGN] OR txid94922[ORGN] OR txid117914[ORGN] OR txid30469[ORGN] OR txid118158[ORGN] OR txid412631[ORGN] OR txid390325[ORGN] OR txid31034[ORGN] OR txid88680[ORGN] OR txid215389[ORGN] OR txid75380[ORGN] OR txid365061[ORGN] OR txid173245[ORGN] OR txid7832[ORGN] OR txid86197[ORGN] OR txid7926[ORGN] OR txid181462[ORGN] OR txid43062[ORGN] OR txid215365[ORGN] OR txid75381[ORGN] OR txid223796[ORGN] OR txid117921[ORGN] OR txid630724[ORGN] OR txid55115[ORGN] OR txid172143[ORGN] OR txid76072[ORGN] OR txid206152[ORGN] OR txid1545873[ORGN] OR txid130264[ORGN] OR txid27767[ORGN] OR txid2748623[ORGN] OR txid274463[ORGN] OR txid242956[ORGN] OR txid172114[ORGN] OR txid30507[ORGN] OR txid603452[ORGN] OR txid443779[ORGN] OR txid2800422[ORGN] OR txid178224[ORGN] OR txid63826[ORGN] OR txid8220[ORGN] OR txid30764[ORGN] OR txid48439[ORGN] OR txid2025516[ORGN] OR txid31099[ORGN] OR txid86359[ORGN] OR txid30840[ORGN] OR txid143897[ORGN] OR txid376643[ORGN] OR txid88694[ORGN] OR txid40580[ORGN] OR txid7790[ORGN] OR txid7855[ORGN] OR txid117842[ORGN] OR txid242959[ORGN] OR txid47697[ORGN] OR txid30986[ORGN] OR txid390339[ORGN] OR txid117889[ORGN] OR txid760215[ORGN] OR txid172118[ORGN] OR txid1489917[ORGN] OR txid27768[ORGN] OR txid30842[ORGN] OR txid215393[ORGN] OR txid30843[ORGN] OR txid8247[ORGN] OR txid30845[ORGN] OR txid81368[ORGN] OR txid8184[ORGN] OR txid97158[ORGN] OR txid30846[ORGN] OR txid130262[ORGN] OR txid30847[ORGN] OR txid390344[ORGN] OR txid30848[ORGN] OR txid242963[ORGN] OR txid183715[ORGN] OR txid30849[ORGN] OR txid8071[ORGN] OR txid81371[ORGN] OR txid30850[ORGN] OR txid30761[ORGN] OR txid8061[ORGN] OR txid30851[ORGN] OR txid57976[ORGN] OR txid7930[ORGN] OR txid88706[ORGN] OR txid181423[ORGN] OR txid30758[ORGN] OR txid30700[ORGN] OR txid30852[ORGN] OR txid8061[ORGN] OR txid170197[ORGN] OR txid908954[ORGN] OR txid57977[ORGN] OR txid31029[ORGN] OR txid143346[ORGN] OR txid88673[ORGN] OR txid30853[ORGN] OR txid30759[ORGN] OR txid118143[ORGN] OR txid8189[ORGN] OR txid30854[ORGN] OR txid7944[ORGN] OR txid46660[ORGN] OR txid68515[ORGN] OR txid1158979[ORGN] OR txid7762[ORGN] OR txid40665[ORGN] OR txid118167[ORGN] OR txid30857[ORGN] OR txid412628[ORGN] OR txid88699[ORGN] OR txid274689[ORGN] OR txid118175[ORGN] OR txid215399[ORGN] OR txid55111[ORGN] OR txid172126[ORGN] OR txid241355[ORGN] OR txid7802[ORGN] OR txid215327[ORGN] OR txid242930[ORGN] OR txid54912[ORGN] OR txid94930[ORGN] OR txid170204[ORGN] OR txid270589[ORGN] OR txid30858[ORGN] OR txid168096[ORGN] OR txid31101[ORGN] OR txid27723[ORGN] OR txid31025[ORGN] OR txid428447[ORGN] OR txid117915[ORGN] OR txid172138[ORGN] OR txid171414[ORGN] OR txid1213740[ORGN] OR txid509847[ORGN] OR txid31102[ORGN] OR txid178226[ORGN] OR txid215337[ORGN] OR txid30859[ORGN] OR txid30860[ORGN] OR txid1545905[ORGN] OR txid8162[ORGN] OR txid178231[ORGN] OR txid390366[ORGN] OR txid210573[ORGN] OR txid206137[ORGN] OR txid30987[ORGN] OR txid170200[ORGN] OR txid117840[ORGN] OR txid30861[ORGN] OR txid8256[ORGN] OR txid30997[ORGN] OR txid490375[ORGN] OR txid81383[ORGN] OR txid30917[ORGN] OR txid112726[ORGN] OR txid30862[ORGN] OR txid30863[ORGN] OR txid30864[ORGN] OR txid30865[ORGN] OR txid55129[ORGN] OR txid260504[ORGN] OR txid55133[ORGN] OR txid30949[ORGN] OR txid1490017[ORGN] OR txid57978[ORGN] OR txid30866[ORGN] OR txid160493[ORGN] OR txid172661[ORGN] OR txid172135[ORGN] OR txid30976[ORGN] OR txid30867[ORGN] OR txid30475[ORGN] OR txid81387[ORGN] OR txid130260[ORGN] OR txid259919[ORGN] OR txid117884[ORGN] OR txid7860[ORGN] OR txid134992[ORGN] OR txid490374[ORGN] OR txid88709[ORGN] OR txid136488[ORGN] OR txid195629[ORGN] OR txid8247[ORGN] OR txid30869[ORGN] OR txid30870[ORGN] OR txid94935[ORGN] OR txid8224[ORGN] OR txid30885[ORGN] OR txid172121[ORGN] OR txid8108[ORGN] OR txid7826[ORGN] OR txid274692[ORGN] OR txid30871[ORGN] OR txid87098[ORGN] OR txid274712[ORGN] OR txid75382[ORGN] OR txid30874[ORGN] OR txid30948[ORGN] OR txid270524[ORGN] OR txid117916[ORGN] OR txid8169[ORGN] OR txid27769[ORGN] OR txid376648[ORGN] OR txid30511[ORGN] OR txid55140[ORGN] OR txid378069[ORGN] OR txid473342[ORGN] OR txid68500[ORGN] OR txid123351[ORGN] OR txid722446[ORGN] OR txid274706[ORGN] OR txid118152[ORGN] OR txid43698[ORGN] OR txid72045[ORGN] OR txid81390[ORGN] OR txid30875[ORGN] OR txid242970[ORGN] OR txid316148[ORGN] OR txid31031[ORGN] OR txid210577[ORGN] OR txid1489881[ORGN] OR txid30480[ORGN] OR txid30876[ORGN] OR txid96776[ORGN] OR txid81393[ORGN] OR txid31027[ORGN] OR txid31026[ORGN] OR txid7839[ORGN] OR txid27770[ORGN] OR txid270659[ORGN] OR txid27774[ORGN] OR txid300225[ORGN] OR txid57771[ORGN] OR txid70846[ORGN] OR txid40662[ORGN] OR txid81374[ORGN] OR txid8243[ORGN] OR txid75383[ORGN] OR txid31103[ORGN] OR txid1489918[ORGN] OR txid8193[ORGN] OR txid56259[ORGN] OR txid42164[ORGN] OR txid28683[ORGN] OR txid8830[ORGN] OR txid8899[ORGN] OR txid30558[ORGN] OR txid9765[ORGN] OR txid85103[ORGN] OR txid8907[ORGN] OR txid8465[ORGN] OR txid8493[ORGN] OR txid9726[ORGN] OR txid27792[ORGN] OR txid37067[ORGN] OR txid29135[ORGN] OR txid8602[ORGN] OR txid30443[ORGN] OR txid43314[ORGN] OR txid37576[ORGN] OR txid184169[ORGN] OR txid85545[ORGN] OR txid8910[ORGN] OR txid226940[ORGN] OR txid9702[ORGN] OR txid56260[ORGN] OR txid30444[ORGN] OR txid37076[ORGN] OR txid30445[ORGN] OR txid9206[ORGN] OR txid9709[ORGN] OR txid9740[ORGN] OR txid9750[ORGN] OR txid30448[ORGN] OR txid30451[ORGN] OR txid425629[ORGN] OR txid66056[ORGN] OR txid8917[ORGN] OR txid9231[ORGN] OR txid54049[ORGN] OR txid30446[ORGN] OR txid33574[ORGN]) AND ("cytochrome c oxidase 1"[Title] OR "cytochrome oxidase subunit I"[Title] OR COI[Title] OR COXI[Title] OR COX1[Title] OR "COX 1"[Title] OR "COX I"[Title] OR CO1[Title] OR C01[Title] OR "cytochrome oxidase I"[Title] OR "cytochrome oxidase subunit I"[Title] OR "cytochrome oxidase subunit 1"[Title] OR "cytochrome oxidase 1"[Title] OR "cytochrome c oxidase subunit I"[Title] OR "cytochrome c oxidase subunit 1"[Title])' | efetch -format fasta > c01_fish_nuccore.fasta

esearch -db gene -query "(txid29146[ORGN] OR txid107764[ORGN] OR txid54907[ORGN] OR txid134615[ORGN] OR txid7850[ORGN] OR txid1489899[ORGN] OR txid205120[ORGN] OR txid84619[ORGN] OR txid1072510[ORGN] OR txid7934[ORGN] OR txid8279[ORGN] OR txid83881[ORGN] OR txid303732[ORGN] OR txid170194[ORGN] OR txid117867[ORGN] OR txid31017[ORGN] OR txid316123[ORGN] OR txid163127[ORGN] OR txid143304[ORGN] OR txid69128[ORGN] OR txid81361[ORGN] OR txid150446[ORGN] OR txid31024[ORGN] OR txid490307[ORGN] OR txid88658[ORGN] OR txid463601[ORGN] OR txid1489799[ORGN] OR txid8065[ORGN] OR txid94935[ORGN] OR txid88661[ORGN] OR txid56718[ORGN] OR txid8253[ORGN] OR txid36203[ORGN] OR txid215350[ORGN] OR txid30757[ORGN] OR txid181401[ORGN] OR txid30850[ORGN] OR txid270592[ORGN] OR txid30908[ORGN] OR txid7865[ORGN] OR txid8157[ORGN] OR txid94930[ORGN] OR txid7805[ORGN] OR txid7802[ORGN] OR txid181420[ORGN] OR txid206122[ORGN] OR txid163122[ORGN] OR txid270595[ORGN] OR txid206106[ORGN] OR txid88666[ORGN] OR txid30828[ORGN] OR txid270610[ORGN] OR txid29142[ORGN] OR txid181415[ORGN] OR txid206140[ORGN] OR txid7869[ORGN] OR txid94304[ORGN] OR txid27583[ORGN] OR txid118146[ORGN] OR txid68511[ORGN] OR txid76914[ORGN] OR txid30943[ORGN] OR txid55118[ORGN] OR txid391217[ORGN] OR txid7941[ORGN] OR txid27766[ORGN] OR txid8092[ORGN] OR txid556237[ORGN] OR txid30947[ORGN] OR txid94922[ORGN] OR txid117914[ORGN] OR txid30469[ORGN] OR txid118158[ORGN] OR txid412631[ORGN] OR txid31034[ORGN] OR txid88680[ORGN] OR txid215389[ORGN] OR txid75380[ORGN] OR txid365061[ORGN] OR txid173245[ORGN] OR txid86197[ORGN] OR txid7926[ORGN] OR txid181462[ORGN] OR txid43062[ORGN] OR txid215365[ORGN] OR txid75381[ORGN] OR txid223796[ORGN] OR txid117921[ORGN] OR txid55115[ORGN] OR txid76072[ORGN] OR txid206152[ORGN] OR txid130264[ORGN] OR txid27767[ORGN] OR txid2748623[ORGN] OR txid274463[ORGN] OR txid242956[ORGN] OR txid30507[ORGN] OR txid603452[ORGN] OR txid63826[ORGN] OR txid8220[ORGN] OR txid30764[ORGN] OR txid48439[ORGN] OR txid86359[ORGN] OR txid30840[ORGN] OR txid143897[ORGN] OR txid376643[ORGN] OR txid88694[ORGN] OR txid40580[ORGN] OR txid7790[ORGN] OR txid7855[ORGN] OR txid242959[ORGN] OR txid47697[ORGN] OR txid30986[ORGN] OR txid390339[ORGN] OR txid1489917[ORGN] OR txid27768[ORGN] OR txid30842[ORGN] OR txid215393[ORGN] OR txid30843[ORGN] OR txid8247[ORGN] OR txid30845[ORGN] OR txid81368[ORGN] OR txid8184[ORGN] OR txid30846[ORGN] OR txid30848[ORGN] OR txid242963[ORGN] OR txid183715[ORGN] OR txid30849[ORGN] OR txid8071[ORGN] OR txid30850[ORGN] OR txid30761[ORGN] OR txid8061[ORGN] OR txid30851[ORGN] OR txid57976[ORGN] OR txid7930[ORGN] OR txid88706[ORGN] OR txid181423[ORGN] OR txid30700[ORGN] OR txid30852[ORGN] OR txid8061[ORGN] OR txid170197[ORGN] OR txid57977[ORGN] OR txid31029[ORGN] OR txid143346[ORGN] OR txid88673[ORGN] OR txid30853[ORGN] OR txid30759[ORGN] OR txid118143[ORGN] OR txid8189[ORGN] OR txid30854[ORGN] OR txid7944[ORGN] OR txid46660[ORGN] OR txid68515[ORGN] OR txid1158979[ORGN] OR txid7762[ORGN] OR txid40665[ORGN] OR txid118167[ORGN] OR txid30857[ORGN] OR txid412628[ORGN] OR txid88699[ORGN] OR txid118175[ORGN] OR txid215399[ORGN] OR txid55111[ORGN] OR txid241355[ORGN] OR txid7802[ORGN] OR txid215327[ORGN] OR txid242930[ORGN] OR txid54912[ORGN] OR txid94930[ORGN] OR txid170204[ORGN] OR txid270589[ORGN] OR txid30858[ORGN] OR txid168096[ORGN] OR txid31101[ORGN] OR txid27723[ORGN] OR txid31025[ORGN] OR txid172138[ORGN] OR txid171414[ORGN] OR txid31102[ORGN] OR txid215337[ORGN] OR txid30859[ORGN] OR txid30860[ORGN] OR txid1545905[ORGN] OR txid8162[ORGN] OR txid178231[ORGN] OR txid30987[ORGN] OR txid170200[ORGN] OR txid117840[ORGN] OR txid30861[ORGN] OR txid8256[ORGN] OR txid30997[ORGN] OR txid81383[ORGN] OR txid30917[ORGN] OR txid30862[ORGN] OR txid30863[ORGN] OR txid30864[ORGN] OR txid30865[ORGN] OR txid55129[ORGN] OR txid260504[ORGN] OR txid55133[ORGN] OR txid30949[ORGN] OR txid57978[ORGN] OR txid30866[ORGN] OR txid172661[ORGN] OR txid30976[ORGN] OR txid30867[ORGN] OR txid30475[ORGN] OR txid130260[ORGN] OR txid259919[ORGN] OR txid117884[ORGN] OR txid7860[ORGN] OR txid134992[ORGN] OR txid490374[ORGN] OR txid88709[ORGN] OR txid136488[ORGN] OR txid195629[ORGN] OR txid8247[ORGN] OR txid30869[ORGN] OR txid30870[ORGN] OR txid94935[ORGN] OR txid8224[ORGN] OR txid8108[ORGN] OR txid7826[ORGN] OR txid274692[ORGN] OR txid30871[ORGN] OR txid87098[ORGN] OR txid75382[ORGN] OR txid30874[ORGN] OR txid30948[ORGN] OR txid270524[ORGN] OR txid117916[ORGN] OR txid8169[ORGN] OR txid27769[ORGN] OR txid376648[ORGN] OR txid30511[ORGN] OR txid55140[ORGN] OR txid378069[ORGN] OR txid68500[ORGN] OR txid123351[ORGN] OR txid274706[ORGN] OR txid118152[ORGN] OR txid43698[ORGN] OR txid72045[ORGN] OR txid81390[ORGN] OR txid30875[ORGN] OR txid242970[ORGN] OR txid316148[ORGN] OR txid31031[ORGN] OR txid210577[ORGN] OR txid30480[ORGN] OR txid30876[ORGN] OR txid96776[ORGN] OR txid81393[ORGN] OR txid31027[ORGN] OR txid31026[ORGN] OR txid7839[ORGN] OR txid27770[ORGN] OR txid270659[ORGN] OR txid27774[ORGN] OR txid300225[ORGN] OR txid70846[ORGN] OR txid8243[ORGN] OR txid75383[ORGN] OR txid31103[ORGN] OR txid1489918[ORGN] OR txid8193[ORGN] OR txid56259[ORGN] OR txid42164[ORGN] OR txid28683[ORGN] OR txid8830[ORGN] OR txid8899[ORGN] OR txid30558[ORGN] OR txid9765[ORGN] OR txid85103[ORGN] OR txid8907[ORGN] OR txid8465[ORGN] OR txid8493[ORGN] OR txid9726[ORGN] OR txid27792[ORGN] OR txid37067[ORGN] OR txid29135[ORGN] OR txid8602[ORGN] OR txid30443[ORGN] OR txid37576[ORGN] OR txid184169[ORGN] OR txid85545[ORGN] OR txid8910[ORGN] OR txid226940[ORGN] OR txid9702[ORGN] OR txid56260[ORGN] OR txid30444[ORGN] OR txid37076[ORGN] OR txid30445[ORGN] OR txid9206[ORGN] OR txid9709[ORGN] OR txid9740[ORGN] OR txid9750[ORGN] OR txid30448[ORGN] OR txid30451[ORGN] OR txid425629[ORGN] OR txid66056[ORGN] OR txid8917[ORGN] OR txid9231[ORGN] OR txid54049[ORGN] OR txid30446[ORGN] OR txid33574[ORGN]) AND ("cytochrome c oxidase 1"[Title] OR "cytochrome oxidase subunit I"[Title] OR COI[Title] OR COXI[Title] OR COX1[Title] OR "COX 1"[Title] OR "COX I"[Title] OR CO1[Title] OR C01[Title] OR "cytochrome oxidase I"[Title] OR "cytochrome oxidase subunit I"[Title] OR "cytochrome oxidase subunit 1"[Title] OR "cytochrome oxidase 1"[Title] OR "cytochrome c oxidase subunit I"[Title] OR "cytochrome c oxidase subunit 1") NOT environmental sample[Title] NOT environmental samples[Title] NOT environmental[Title] NOT uncultured[Title] NOT unclassified[Title] NOT unidentified[Title] NOT unverified[Title] NOT "cytochrome b"[Title] NOT chromosome[Title] NOT "cytochrome P450"[Title] " | efetch -format docsum | xtract -pattern GenomicInfoType -element ChrAccVer ChrStart ChrStop | while IFS=$'\t' read acn str stp; do efetch -db nuccore -format fasta -id "$acn" -chr_start "$str" -chr_stop "$stp"; done | sed "s/>/>16S_/" > c01_fish_gene.fasta
